# Storage Protein-Mediated Translation Control Links Juvenile Diet to Longevity in *Drosophila*

**DOI:** 10.1101/2025.05.18.654697

**Authors:** Hina Kosakamoto, Rina Okada, Clive Barker, Jun Seita, Naoshi Dohmae, Koshi Imami, Fumiaki Obata

## Abstract

Dietary restriction (DR), whether applied during adulthood or juvenile stages, extends lifespan across diverse species. However, the mechanisms by which early-life dietary interventions influence adult physiology and longevity remain poorly understood. Here, using *Drosophila* as a model, we demonstrate that protein restriction during the larval stage (early-life protein restriction, ePR) promotes adult lifespan by reducing storage protein levels. Stable isotope tracing reveals that dietary amino acids obtained in the larval stage are retained into adulthood, especially incorporated into ribosomal proteins. This is mediated by larval serum protein 2 (Lsp2), a major storage protein, whose expression is durably downregulated by ePR. Both dietary (ePR) and genetic (Lsp2-RNAi) reduction of the protein storage lead to decreased ribosomal protein levels and translation activity in early adulthood. Notably, these storage proteins are enriched in aromatic amino acids such as tyrosine, and larval tyrosine restriction alone is sufficient to suppress translation and promote longevity. These findings show that storage proteins mediate the effect of larval nutrition on adult longevity via controlling translation. Our study uncovers a previously unrecognised mechanism of nutritional memory that links early-life diet to life-long organismal health.

## Introduction

Nearly a century ago, dietary restriction (DR), without malnutrition, was shown to delay ageing and extend lifespan in rats^1^. Since then, research in the DR-longevity field has primarily focused on adult-onset interventions, in both young and aged animals, as these are more translatable to potential anti-ageing strategies in humans. However, early foundational studies also implied that restricting diet exclusively during early-life periods was sufficient to extend lifespan in rats^1,2^. This phenomenon of “early-life DR” was reproduced in the invertebrate *Daphnia longispina*^3^ and subsequently observed in other model organisms, including the fruit fly *Drosophila melanogaster*^4–6^ and mice^7–9^. Several of these studies suggest that restricting protein alone during juvenile stages— such as lactation and growth— can be sufficient to extend lifespan. Similarly, early-life administration of rapamycin—an inhibitor of the amino acid sensing mechanistic Target of Rapamycin complex 1 (mTORC1)—has been shown to promote longevity in both flies and mice^10,11^. Despite consistent evidence across species, the molecular mechanisms by which early-life nutritional cues are retained and translated into long-term physiological changes remain poorly understood. In particular, the identity of molecules that encode, maintain, and transmit this juvenile nutritional memory into adult is yet to be fully elucidated.

To dissect the long-term physiological consequences of early-life nutrient availability, it is critical to employ genetic tools capable of manipulating candidate pathways during development and assessing their effects throughout life. However, such investigations are technically challenging due to the complicated nature of the phenomenon and time-consuming longitudinal analysis of ageing and longevity. In this context, *Drosophila* offers a powerful system: its lifecycle comprises well-defined developmental transitions from embryo to adult in just ten days, and its short lifespan facilitates rapid lifespan analysis in a few months. Furthermore, advanced genetic and dietary tools enable precise spatial and temporal control of gene function and nutritional response. In this study, we examined how protein restriction during the juvenile stages impacts adult physiology and longevity, and identified storage proteins as key mediators linking early-life nutrition to adult health outcome.

## Results

### Early-life protein restriction (ePR) extends lifespan

Late second-instar larvae were transferred to a low yeast diet at approximately 68 hours after egg laying (AEL), and after eclosion, the adult flies were returned to a standard yeast-based diet (Fig. 1a). Since yeast is a major source of protein, this leads to the transient, larval protein restriction. We first assessed the effect of the low protein diet on developmental speed, eclosion rate, and body growth (Fig. 1b-e). Reducing yeast content in the larval diet from 8% to 2% or 1% led to a delayed developmental speed (Fig. 1b). Once larvae reached the pupal stage, the eclosion rate remained unaffected (Fig. 1c). Protein restriction decreased adult body weight in a dose-dependent manner in both females and males (Fig. 1d,e). Female flies subjected to protein restriction during larval stages exhibited up to a 28% reduction in body weight (Fig. 1d,f). The size of wings, a hallmark of body size, was also decreased (Fig. 1g). In addition to the smaller body size, we noticed that the body wall displayed a slightly lighter coloration especially at the abdomen (Fig. 1f), suggestive of inadequate pigmentation derived from nutrient including an amino acid tyrosine (Tyr)^12–14^.

**Fig. 1:**
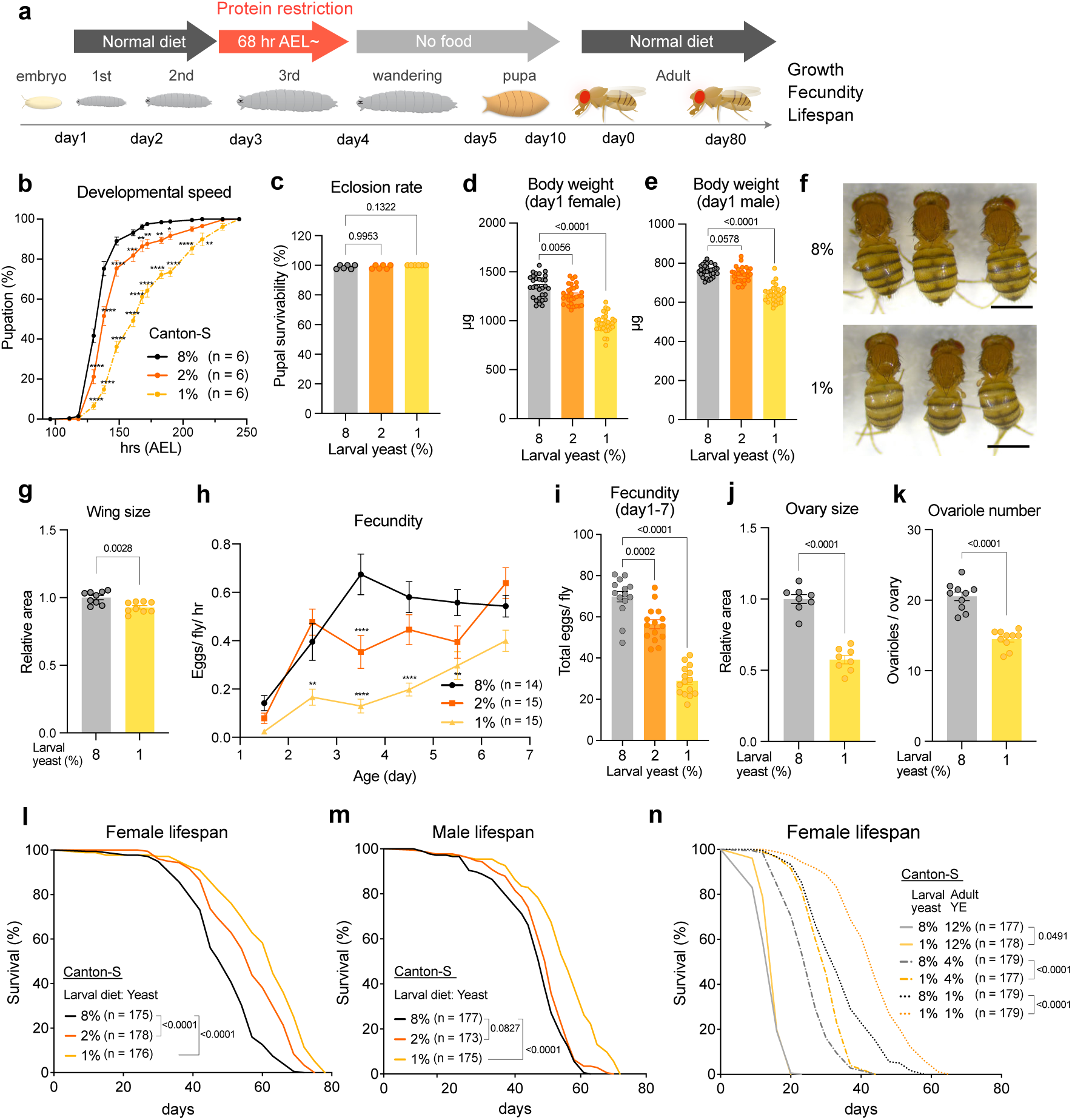
Larval protein restriction impacts adult physiologies and extends lifespan. **a**, Experimental scheme of early-life protein restriction (ePR). **b**, Developmental speed of Canton-S flies with ePR. n = 6. **c**, Eclosion rate of Canton-S flies with ePR. n = 6. **d**,**e**, Body weight of Canton-S female (**d**) and male (**e**) flies with ePR at day1 after eclosion. n = 30. **f**, Images of the Canton-S female flies with ePR at day1 after eclosion. Scale bar = 1 mm. **g**, Wing size of Canton-S female flies with ePR. n = 9. **h**,**i**, Fecundity per day (**h**) or per week (**i**) of Canton-S female flies with ePR from day1 to day7 after eclosion. (n = 14 (8%) or 15 (2% and 1%). **j**,**k**, Ovary size (i) and ovariole number of Canton-S flies with ePR at day4. n = 8 (**j**) or 10 (**k**). **l**,**m**, Lifespan of Canton-S female (**l**) or male (**m**) flies with ePR. Sample sizes (n) are shown in the figure. **n**, Lifespan of Canton-S female flies with ePR and adult feeding of various protein diet. Sample sizes (n) are shown in the figure. For the statistics, two-way ANOVA with Dunnett’s multiple comparison test (**a**,**h**), one-way ANOVA with Dunnett’s multiple comparison test (**c-e, i**), a two-tailed Student’s *t* test (**g,j,k**), or a log-rank test (**l**-**n**) was used. \**P* < 0.05; \*\**P* < 0.01; \*\*\**P* < 0.001; \*\*\*\**P* < 0.0001. For all graphs, the mean and SEM are shown. Data points indicate biological replicates.

Next, we assessed for phenotypes of the adult flies after this early-life protein restriction (ePR). Although reared on a standard diet at this stage, adult female flies showed compromised fecundity, particularly during the early adult stage (Fig. 1h,i) with concomitant observation of the reduced ovary size and a decreased number of ovarioles (Fig. 1j,k). The number of ovarioles is determined during the larval stage and is affected by larval nutrition^15^. These results suggest that ePR induces irreversible developmental effects on body growth and the reproductive organ. Given that the decrease in the ovariole number is milder than the one observed in the fecundity, additional mechanism is likely to influence the egg production.

Consistent with the previous findings^6^, adult flies with ePR exhibited an increased lifespan in both females and male flies (Fig. 1l,m). Notably, the lifespan-prolonging effect of ePR seemed to be independent from the adult protein restriction as the lifespan extension by ePR is additive to the adult manipulation (Fig. 1n). In contrast, lifespan extension by ePR was attenuated when flies were provided with a high-protein diet in adulthood (Fig. 1n). We then tested how their lifespan was affected by various concentration of dietary amino acids at the adult stage using a synthetic diet (Extended Data Fig. 1a-g)^16,17^. The lifespan extension by ePR was observed within the range of 10%-100% amino acids (Extended Data Fig. 1a-d). Moreover, the lifespan extension under moderate AA restriction (10%-50%) was consistently observed regardless of ePR (Extended Data Fig. 1f,g), suggesting that the mechanisms underlying lifespan extension by ePR differ from those of adult protein restriction. Interestingly, however, the lifespan of adult flies with ePR was compromised at 0% amino acid (Extended Data Fig. 1e). Thus, inadequate nutrient during the larval stage is detrimental when adult flies have no access to dietary amino acids, and therefore, ePR is not universally beneficial. It also suggests that the effect of ePR is modulated by adult dietary conditions and therefore the outcome of ePR can be variable among experimental settings^6,18^.

### ePR alters nutrient signalling in early adulthood

Lifespan extension is closely linked to downregulation of insulin signaling^19^. We found that ePR reduced phospho-Akt levels in the abdominal carcass (enriched in the fat body) of day2 adult female flies, indicating suppressed insulin signalling (Extended Data Fig. 2a,b). Consistent with this, activation of the transcription factor Forkhead Box, sub-group O (FoxO), which is typically repressed by insulin signalling, was suggested by the upregulation of its target genes, including *insulin-like receptor* (*InR*) and *insulin-like peptide 6* (*Ilp6*), in the abdominal carcass (Extended Data Fig. 2c,d). Expression of *female-specific independent of transformer* (*fit*) visualised by a GFP reporter was also suppressed in the abdomen by ePR, reflecting reduction of insulin/TOR signalling activity (Extended Data Fig. 2e)^20,21^. Decreased expression of insulin-like peptides in the head further corroborated the suppression of insulin signalling activity (Extended Data Fig. 2f-h). However, this suppression was transient, as phospho-Akt levels and *InR* expression returned to control levels at day 7 after eclosion (Extended Data Fig. 2i-k).

In our previous study, we found that protein restriction activates the amino acid sensing pathway, activating transcription factor 4 (ATF4), in the larval fat body^22^. To ask whether the ATF4 activation is persistent, we visualised the ATF4 activity by using *4E-BP^intron^-dsRed* reporter line^23^. ATF4 was activated by ePR until day 2 of adulthood, but this returned to baseline at day 5 of adulthood (Extended Data Fig. 2l,m). Despite ATF4 activation, the upstream phospho-eIF2α levels was not altered in the abdominal carcass of day 2 flies, suggesting eIF2α-independent ATF4 activation (Extended Data Fig. 2n,o). Taken together, nutrient restriction in the larval stage affects nutrient sensing pathways transiently in early adulthood.

### Adult proteins are derived from larval diet

Our phenotypic analysis suggested that the effect of early-life protein intake was persistent throughout larval, pupal, and early adult stages. We hypothesised that some proteins or amino acids during larval stage remains to adulthood. To trace dietary amino acids, we employed *in vivo* pulse SILAC (pSILAC) by using synthetic diet. We fed larvae with the synthetic diet which contains “heavy” isotope-labelled lysine (Lys) and arginine (Arg), instead of “light (natural)” ones (Fig. 2a). This enabled to trace the dietary amino acid fate by entirely swapping the two amino acids with the stable isotopes without disturbing their physiology and developmental process. After eclosion, the adult flies were reared on the standard yeast-based diet containing “light” amino acids for two days and then switched to a synthetic diet containing “medium” isotope-labelled Lys and Arg. Following four days of feeding the synthetic diet in the adult, we quantified the isotopic composition of each protein in the head (contained not only the brain but also many other tissues including the fat body) and the abdominal carcass (contained the fat body enriched) using nano-liquid chromatography and tandem mass spectrometry (nanoLC/MS/MS). The duration of isotope labelling (four days) in the larval stage (heavy isotope) and the adult stage (medium isotope) were matched to enable direct comparison.

**Fig. 2:**
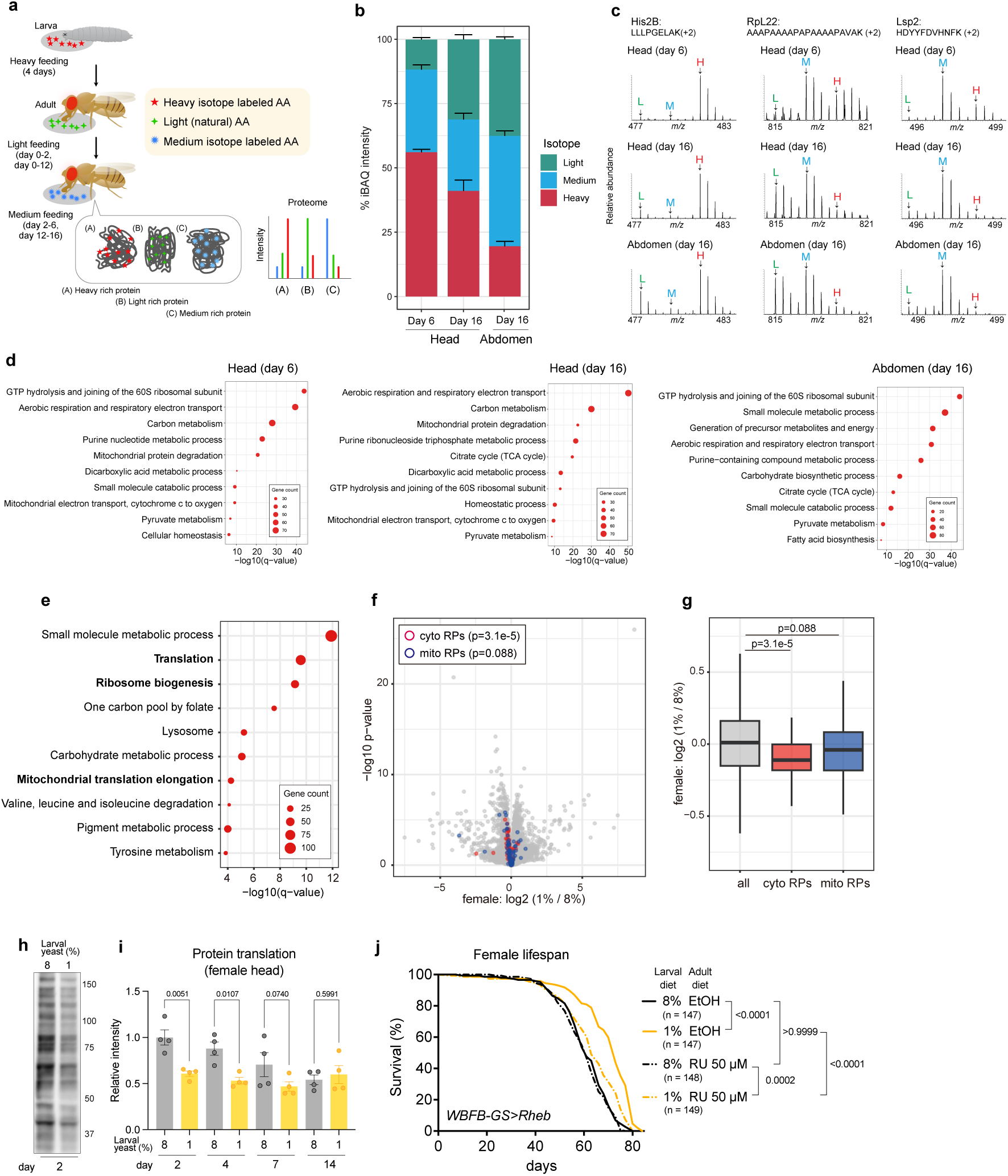
Stable isotope labelling and proteome analysis of flies after ePR. **a**, Experimental scheme of the *in vivo* pulse SILAC analysis. Heavy isotope-labelled amino acids were provided during the larval stage and following eclosion standard yeast diet were provided for either 2 days or 12 days. Then, medium isotope-labelled amino acids were provided for 4 days. **b**, Proportions of abundance of light-, medium-, and heavy-labelled proteins within the head and abdomen samples of Canton-S female flies. Intensity-based absolute quantification (iBAQ) algorithm computes the sum of all the peptides intensities divided by the number of theoretically observable peptides, which provides rough estimation of protein abundance. **c**, Exemplary MS1 spectra for three peptides derived from His2b (slow turnover), RpL22 (intermediate turnover), and Lsp2 (fast turnover). **d**, GO analyses showing the ten most enriched gene ontology terms of top 25% abundant heavy-labelled proteins. Q-values are computed based on adjusted p-values for multiple testing with Benjamini-Hochberg and −log10-transformed. **e**, GO enrichment analysis of proteins which expression was significantly decreased by ePR in heads of Canton-S female flies at day2 (p<0.01 and log2 1% / 8% <-0.1). **f**, A volcano plot showing mean log2 fold change (1% / 8%) and −log10 p-value of proteins in the head of Canton-S female flies with ePR at day2. The cytosolic and mitochondrial ribosomal proteins are indicated by red and blue circles, respectively. **g**, A box plot showing log2 fold change (1% / 8%) of proteins grouped into three categories: all proteins, cytosolic and mitochondrial ribosomal proteins. **h**, A representative image of protein translation using SUnSET assay in the heads of Canton-S female flies with ePR at day2-3. **i**, Quantification of protein translation using SUnSET assay in the heads Canton-S female flies with ePR during day2-3, 4-5, 7-8, and 14-15. n = 4. **j**, Lifespan of female flies with ePR and *Rheb* overexpression using adult fat body driver (WBFB-GS). Food supplemented with either RU486 (RU) or 100% ethanol (EtOH) was provided from day0. Sample sizes (n) are shown in the figure. For the statistics, two-sided Wilcoxon rank-sum test (**g**) or one-way ANOVA with Holm-Šídák’s multiple comparison test (**i**) was used. For the graphs, the mean and SEM (**b,i**) or the minimum, the lower quartile, the median, the upper quartile, and the maximum points (**g**) are shown. Data points indicate biological replicates.

From the proteomic analysis, we obtained a reproducible result as revealed by high correlation of H/M ratios among biological replicates (Extended Data Fig. 3a). When comparing amino acids from the larval stage (heavy) with those obtained during adulthood from day 2–6 (medium) in the adult proteome of the head, we observed a general trend that heavy isotope (56.1% of the total) was more abundant than medium-isotope (32.3%) or even medium-plus light-amino acids (Fig. 2b, Supplementary Table 1). This finding indicates that early adult proteome contains a substantial proportion of proteins derived from larval dietary amino acids (Fig. 2c).

Next, we prepared flies with the medium-labelled diet at a later stage, *i.e* day12. In this experimental setup, the flies were fed light (yeast-based) amino acids for 12 days, then fed medium amino acids for four days (Fig. 2a). At this stage (day 16), although the proportion of heavy isotopes were decreased compared to day-6 samples, heavy-labelled proteome (41.0% of the total) remained more prevalent than medium-labelled one (27.8%) (Fig. 2b, Supplementary Table 1). In the abdomen at day 16, the proportion of heavy-labelled proteome is relatively low (19.5% of the total), but substantial amount still remained in the adulthood (Fig. 2b). The difference between head and abdomen is the existence of the brain, which does not undergo histolysis during pupal stage. These findings together suggest that amino acids ingested during the larval stage remain in the adult proteome for more than two weeks. There may be two possibilities to explain this results; 1) the long-lived proteins are carried over from the larval stage or 2) the larva-derived proteins are efficiently recycled to produce adult proteins. Indeed, long-lived proteins such as Histone H2B (His2B) was enriched with heavy isotope-labelled amino acids (Fig. 2c).

Gene ontology enrichment analysis revealed that top 25% abundant proteins labelled by heavy isotopes were predominantly involved in metabolic processes in mitochondria and protein translation (Fig. 2d). We found that 41 ribosomal proteins were enriched in the early adult proteome (Supplementary Table 1). For instance, 60S ribosomal protein L22 (RpL22) exhibited significant amount of heavy isotope labelling at day 6, although the ratio of H/M rapidly goes down at day 16 suggests that RpL22 has higher turnover (Fig. 2c). These data indicates that metabolic and ribosomal proteins, especially at the early adulthood, are carried over from the larval stage or made up from the larva-obtained amino acids. Analysis of incompletely digested chimeric peptides derived from RpL22 revealed the presence of peptides containing both heavy- and middle-labelled lysine in head samples at day 6 (Extended Data Fig. 3b). This finding implies that larval-derived proteins are recycled to synthesize adult proteins at early stages, although the frequency of such chimeras was lower (Extended Data Fig. 3b). In contrast, His2B contains little chimeric peptides labelled with both isotopes, consistent with its known characteristic as a long-lived protein (Extended Data Fig. 3b).

### ePR alters proteome and decreases translation in early adult

Our *in vivo* pSILAC analysis suggested that amino acids obtained during larval stage remains in adulthood and some proteins can be particularly influenced by larval protein restriction. To understand how ePR influenced adult proteome quantitatively, we performed amino acid quantification and proteome analysis of early adult flies upon ePR. We first implemented acid hydrolysis of whole-body samples (see Methods) and quantified the total amino acid content, encompassing both amino acids incorporated into structural and functional proteins as well as free amino acids. ePR led to a significant reduction in total amino acid content, even after normalisation to body weight, both in females and males (Extended Data Fig. 4a,o). Beyond a reduction in total levels, we observed alterations in amino acid composition (Extended Data Fig. 4c,o). For instance, valine, alanine and proline were increased, while Arg, Lys, phenylalanine, and Tyr were decreased by ePR in female flies (Extended Data Fig. 4c). Such compositional changes suggested modifications in the amino acid profile by ePR and existence of a bias in “carry-over” amino acids and proteins.

From the proteome analysis of day 2 adult flies upon ePR (Extended Data Fig. 5a-d), we found that biological processes related to translation, ribosome biogenesis, and mitochondrial translation elongation, as well as metabolic processes, were downregulated in response to ePR (Fig. 2e), which is consistent with our *in vivo* pSILAC data. Indeed, cytosolic and mitochondrial ribosomal proteins were expressed at lower levels in day 2 adult female flies after ePR (Fig. 2f,g). Similar changes were also observed in male flies (Extended Data Fig. 5e-g). These findings led us to hypothesise that translation capacity in early adulthood could be suppressed by ePR. We also noticed that tyrosine and pigment metabolism was decreased (Fig. 2e, Extended Data Fig. 5e), which is consistent with the fact that they are pale (Fig. 1f).

To directly test the impact of larval protein intake on translation in adult flies, we performed a puromycin incorporation assay, which labels newly synthesised peptides. As we expected, protein synthesis was downregulated in the early adult flies upon ePR (Fig. 2h,i). We found that protein synthesis rate was high in early adulthood and gradually decreased during ageing (Fig. 2i). This is consistent with a previous study demonstrating that early-adult spike in translation plays a critical role in determining adult lifespan^24^. The impact of ePR on suppressed translation did not last more than a week (Fig. 2i), however the suppressing translation only in a first week is known to be sufficient for lifespan extension^24^. By enhancing translation by overexpressing *Rheb* (mTORC1 activator) in the adult fat body, the lifespan extension by ePR was attenuated (Fig. 2j). Taken together, our results suggest that larval protein intake influences the early adult translation, even when they are switched back to a standard diet after eclosion, which in turn regulates organismal lifespan.

### Larval protein restriction decreases adult Lsp2 expression

Next, we sought to identify molecules mediating ePR’s effect on adult translation and lifespan. When we conducted transcriptome analysis of the head and abdomen in day 2 adult flies after ePR (Fig. 3a), two storage protein genes, *Larval serum protein 1β* (*Lsp1β*) and *Lsp2*, were among the downregulated genes in the head (Fig. 3b, Extended Data Fig. 6a). The suppression of *Lsp2* was also observed in abdominal samples (Extended Data Fig. 6b). Gene ontology analyses on the up- and down-regulated genes indicate that ePR upregulates cuticle-related genes and stress response pathways in both the head and abdomen (Extended Data Fig. 6c,o). Metabolic genes involved in amino acid, carbohydrate, and nucleotide metabolism were downregulated in the head (Extended Data Fig. 6e,o). These findings suggest that flies exposed to ePR may exhibit adaptive responses to immature cuticle formation and altered protein composition (Fig. 2e-g), while simultaneously downregulating metabolic activity. In addition, there is a tendency of suppression of genes encoding cytosolic ribosomal proteins in flies after ePR (Extended Data Fig. 7a), while mitochondrial ribosomal protein genes remained unaffected (Extended Data Fig. 7b). This is interesting given that both cytosolic and mitochondrial ribosomal proteins were decreased in the proteome analysis. These data suggested that these ribosomal proteins, at least mitochondrial ones, are regulated at the protein level.

**Fig. 3:**
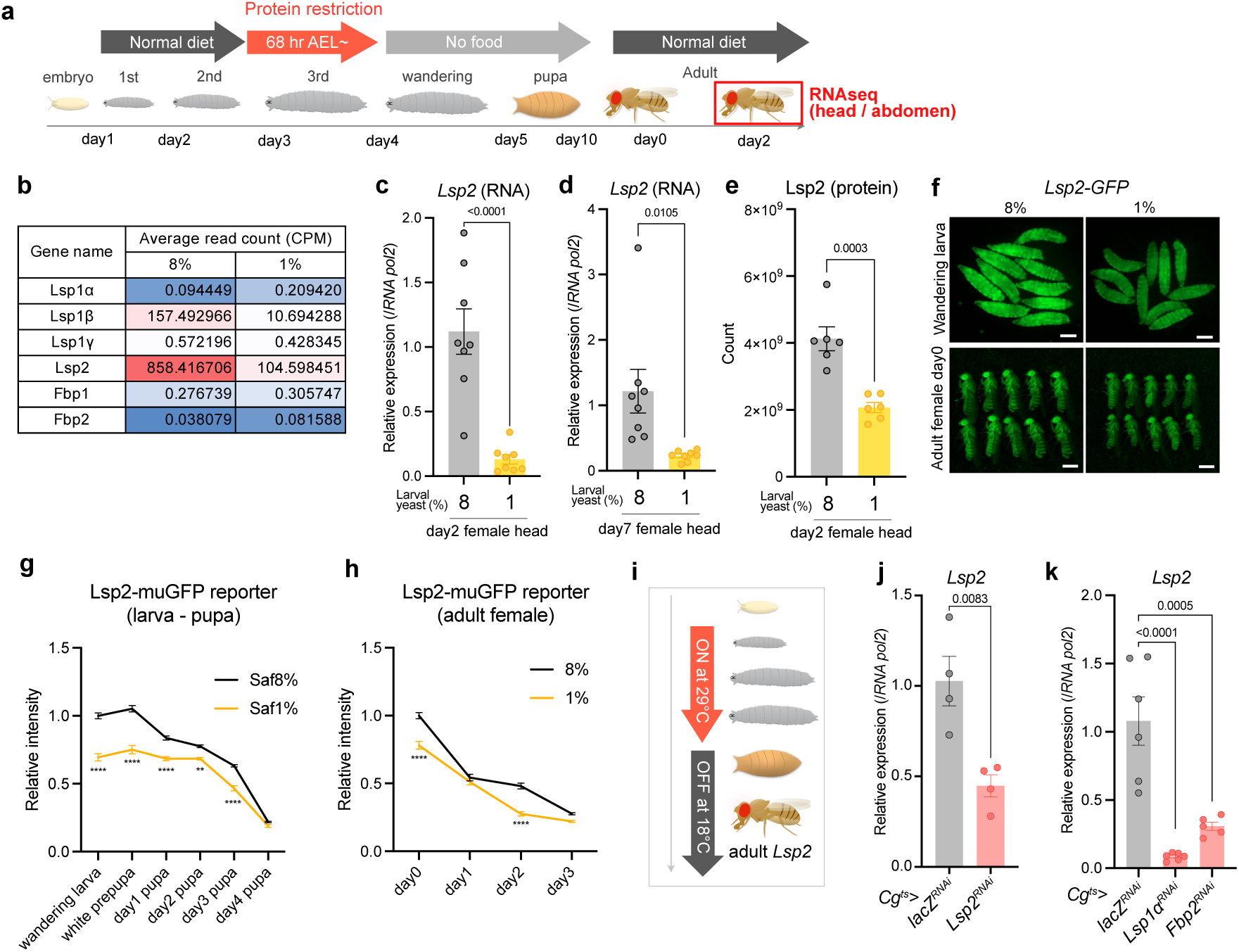
Larval protein restriction suppresses adult *Lsp2* expression. **a**, Experimental scheme for adult transcriptome upon ePR. **b**, Average read count of storage proteins from transcriptome analysis in the Canton-S female flies with ePR at day2. n = 4. **c**,**d**, Quantitative RT‒PCR analysis of *Lsp2* in the Canton-S female flies with ePR at day2 (**c**) or day7 (**d**). n = 8. **e**, Protein expression of Lsp2 from standard proteome analysis in the Canton-S female flies with ePR at day2. **f-h**, Representative images (**f**) and their quantification (**g**,**h**) of the *Lsp2-muGFP* reporter animals with ePR. Scale bar = 1 mm. n = 10 (larvae and adults) or 8 (pupae). **i**-**k**, Experimental scheme (**i**) and quantitative RT-PCR analysis of *Lsp2* after larval-stage specific knockdown of *Lsp2* (**j**) and *Lsp1α* or *Fbp2* (**k**) using fat body driver *Cg-Gal4* with *tub-Gal80^ts^*. n = 4. For the statistics, a two-tailed Student’s *t* test (**c**-**f**, **j**), two-way ANOVA with Šídák’s multiple comparison test (**g**,**h**), or one-way ANOVA with Dunnett’s multiple comparison test (**k**) was used. \*\*\*\**P* < 0.0001. For all graphs, the mean and SEM are shown. Data points indicate biological replicates.

Lsps are known to serve as storage proteins that provide an amino acid reservoir during the pupal (non-feeding) stage for the construction of adult tissues^25^. Lsp proteins are primarily produced in the larval fat body, released into the haemolymph, and subsequently taken up by the fat body during pupal development. *Lsp2* and other storage proteins are highly expressed in the late larvae and early pupal stage, and the expression is dramatically decreased toward the end of pupal development^25^. Nevertheless, our RNA sequencing analysis detected unexpectedly high read counts of *Lsp2* and *Lsp1β* in the heads of adult female flies fed with 8%-yeast diet, whereas other storage proteins showed much lower expression levels (Fig. 3b). Especially, Lsp2 was one of the highly abundant proteins in adult heads found in the proteome analysis (Supplementary Table 2). We performed quantitative RT-PCR and confirmed the downregulation of *Lsp2* expression in female heads by ePR at both day 2 and day 7 (Fig. 3c,d). Proteome analysis and western blotting further confirmed the decreased Lsp2 protein level in day 2 head upon ePR (Fig. 3e, Extended Data Fig. 8a,o).

It is known that larval fat body persists in the adult flies until day2, suggesting that Lsp2 detected in adult flies may originate from this residual tissue^26^. To monitor the Lsp2 protein dynamics in greater detail, we created the *Lsp2-monomeric ultrastable Green Fluorescent Protein* (*muGFP*) knock-in flies by CRISPR/Cas9. As expected, we observed that Lsp2-muGFP protein is abundantly present in the haemolymph in the late larval stage (Fig. 3f). The Lsp2-muGFP levels declined during pupal stage and further decreased in the early adulthood (Fig. 3f-h). By protein restriction in the larval stage, the Lsp2-muGFP intensity was decreased throughout the life stages (Fig. 3f-h).

Given the decrease of adult *Lsp2* mRNA expression and Lsp2-muGFP protein expression in the larval to early adult stage, it was conceivable that *Lsp2* messenger level is kept downregulated from the larval stage with ePR. However, *Lsp2* mRNA expression was not decreased by ePR at the wandering larvae and early pupal stages (Extended Data Fig. 8c). From day 4 pupa onward, the *Lsp2* mRNA levels were again decreased by ePR (Fig. 3c, Extended Data Fig. 8c) Interestingly, when we transiently knocked down *Lsp2* in the larval fat body using the fat body driver *Cg-Gal4* combined with a temperature sensitive form of Gal80, *Lsp2* mRNA expression in early adults was decreased (Fig. 3i,j). These results suggest that the loss of Lsp2 protein during larval stages leads to transcriptional suppression of the *Lsp2* gene in adults. To examine whether this suppression is a general consequence of losing any storage protein, we knocked down other storage proteins—Larval Serum Protein 1α (*Lsp1α*) and Fat Body Protein 2 (*Fbp2*)—using RNAi lines previously verified to lack off-target effects on *Lsp2*^25^. Consequently, we found that knockdown of *Lsp1α* or *Fbp2* during the larval stage also resulted in reduced *Lsp2* expression in adults (Fig. 3k), suggesting that there is a feed forward mechanism to regulate storage protein expression.

These results together suggested the possible mechanism of nutritional memory in ePR: Protein restriction in the larval stages decreases larval storage proteins. Loss of storage protein during larval stage leads to decreased protein and transcription of *Lsp2* in adult stage. Storage proteins are the source of amino acid required to produce new protein, and thus the decrease of their levels decreases protein synthesis, and influence adult proteome. Interestingly, Lsp2 protein exhibits a rapid turnover, as evidenced by the *in vivo* pSILAC experiment, which showed greater enrichment of middle labelling (adult-derived) amino acids compared to heavy (larva-derived) ones (Fig. 2c). This indicates that adult Lsp2 is synthesized *de novo* in the adult stage using dietary amino acids, rather than simply carryover from the larval stage. Therefore, the observed reduction of Lsp2 protein in adult is probably due to the suppression of its transcription.

### Loss of Lsp2 suppresses protein translation and extends lifespan in adulthood

To test functional contribution of Lsp2 for phenotypes observed in the flies after ePR, we knocked down *Lsp2* in the fat body. We found that *Lsp2* knockdown resulted in a lighter body colour especially at the abdominal pigment, similar to the effect observed in ePR (Fig. 4a). *Lsp2* knockdown did not affect adult body weight under sufficient dietary yeast conditions but decreased under a low-protein diet, suggesting that Lsp2 acts as a buffering system to maintain growth under limited protein intake (Extended Data Fig. 9a). *Lsp2* knockdown did not strongly compromise female fecundity (Extended Data Fig. 9b,o) nor the number of ovarioles (Extended Data Fig. 9d). These results suggest that the decrease in body growth and reproduction of ePR is not mediated by Lsp2. In addition, abdominal insulin signalling remained unchanged by *Lsp2* knockdown (Extended Data Fig. 9e-h). Meanwhile, the ATF4 activity was persistently elevated during development in response to *Lsp2* knockdown (Extended Data Fig. 9i,o).

**Fig. 4:**
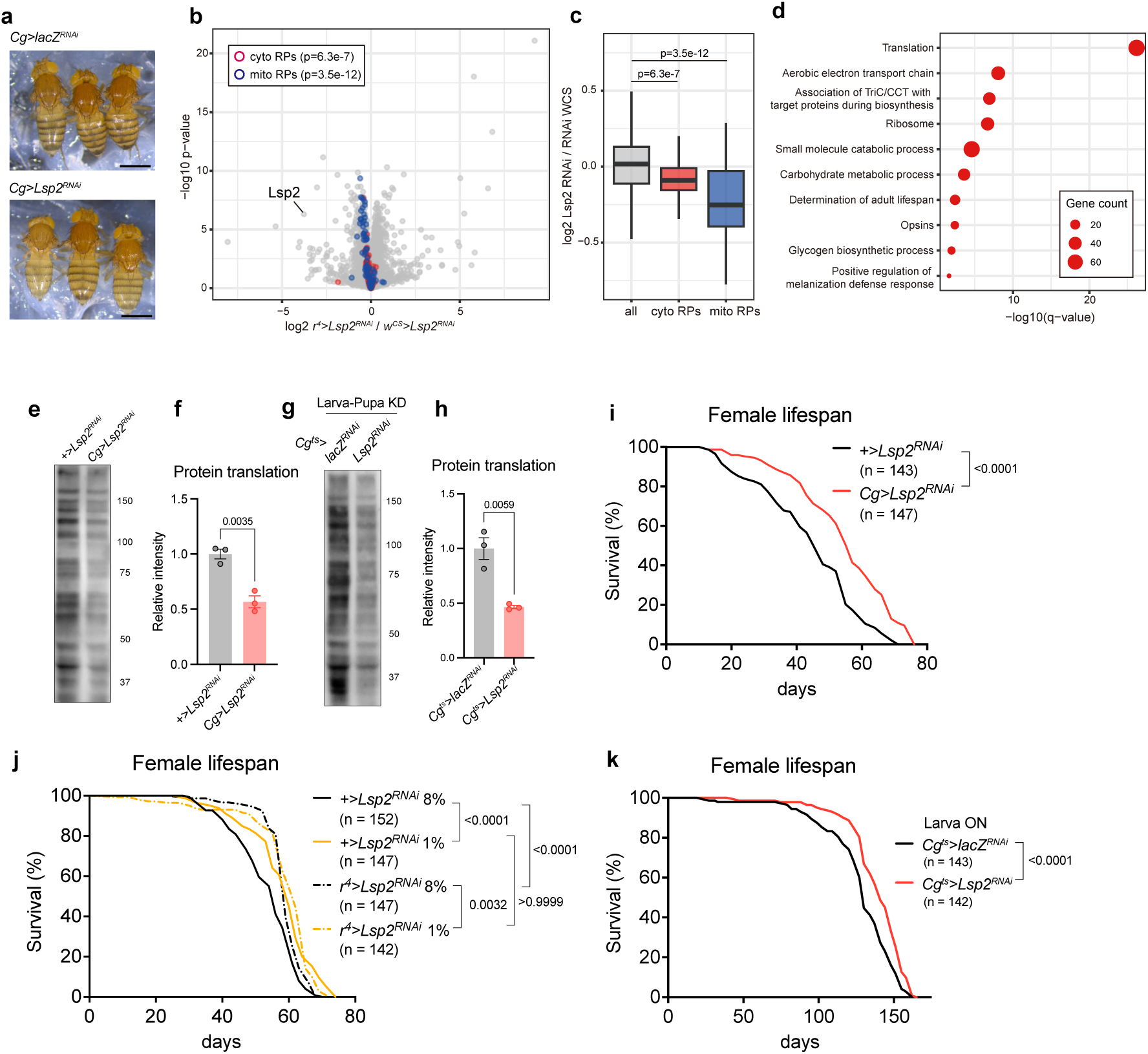
Lsp2-RNAi decreases adult translation and extends lifespan. **a**, Representative images of adult female flies at day0 with *Lsp2* knockdown using fat body driver *Cg-Gal4*. Scale bar = 1 mm. **b**, A volcano plot showing mean log2 fold change (*r^4^>Lsp2^RNAi^* / *w^CS^>Lsp2^RNAi^*) and −log10 p-value of proteins in the head of female flies at day2. The cytosolic and mitochondrial ribosomal proteins are indicated by red and blue circles, respectively. **c**, A box plot showing log2 fold change (*r^4^>Lsp2^RNAi^*/ *w^CS^>Lsp2^RNAi^*) of proteins grouped into three categories: all proteins, cytosolic and mitochondrial ribosomal proteins. **d**, GO enrichment analysis of proteins which expression was significantly decreased by *Lsp2-RNAi* in heads of female flies at day2 (p<0.01 and log2 *r^4^>Lsp2^RNAi^* / *w^CS^>Lsp2^RNAi^*<-0.1). **e**-**h**, A representative image of protein translation analysis (**e**,**g**) and its quantification (**f**,**h**) in the heads of *Lsp2*-knocked down adult flies at day2-3. *Lsp2* knockdown was performed continuously (**e**,**f**) or only during larval and pupal stages (**g**,**h**). Anti-puromycin antibody is used. n = 3. **i**, Female lifespan of *Lsp2*-knocked down flies using fat body driver *Cg-Gal4*. Sample sizes (n) are shown in the figure. **j**, Female lifespan of *Lsp2*-knocked down flies using fat body driver *r^4^-Gal4* with larval protein restriction. Sample sizes (n) are shown in the figure. **k**, Female lifespan of *Lsp2*-knocked down flies using fat body driver *Cg-Gal4* combined with *tub-Gal80^ts^*. The knockdown was performed during larval stage (1^st^ instar larva to wandering larva). Sample sizes (n) are shown in the figure. For the statistics, two-sided Wilcoxon rank-sum test (**c**), a two-tailed Student’s *t* test (**f**,**h**), or a log-rank test (**i**-**k**) was used. For all graphs, the minimum, the lower quartile, the median, the upper quartile, and the maximum points (**c**) or the mean and SEM (**f**,**h**) are shown. Data points indicate biological replicates.

We then performed both transcriptome and proteome analyses of flies with *Lsp2-*RNAi. As expected, we observed a tendency for cytosolic ribosomal protein to be suppressed, whereas mitochondrial ribosomal protein genes remained unaffected (Extended Data Fig. 10a,o). Proteome analysis of *Lsp2* knocked down flies revealed the suppression of both cytosolic and mitochondrial ribosomal proteins, as well as metabolic genes (Fig. 4b-d and Extended Data Fig. 11). These omics results were consistent with those of ePR. Likewise, *Lsp2* knockdown flies had a suppressed translation activity (Fig. 4e,f). Knocking down *Lsp2* specifically during developmental stage also led to the downregulated translation in early adults, demonstrating that the larva-regulated Lsp2 mediates the translation control in adulthood (Fig. 4g,h). Strikingly, knockdown of *Lsp2* extended adult lifespan of female flies (Fig. 4i,j). The lifespan extension by *Lsp2-RNAi* was comparable to that of larval protein restriction (Fig. 4j). ePR failed to further increase lifespan when it combined to *Lsp2-RNAi* (Fig. 4j), suggesting that the two manipulations share the mechanism. *Lsp2* knockdown similarly extended male lifespan (Extended Data Fig. 12a,o). *Lsp2* knockdown only in the larval stage was sufficient to extend lifespan (Fig. 4k). These findings together suggested that Lsp2 plays a central role in mediating the effects of ePR on adult translation and lifespan, but not growth, fecundity, or insulin signalling.

### Tyrosine restriction in larval stage extends lifespan

Lsp2 belongs to the arylphorin class of proteins, which is uniquely enriched in aromatic amino acids. Specifically, Lsp2 protein contains 1.97-fold more Phe and 2.66-fold more Tyr than the average fly protein, i.e. exome^16^ (Fig. 5a). Consequently, knocking down *Lsp2* significantly reduced the overall composition of the two amino acids (Fig. 5b). The amino acid profile of Lsp2-RNAi flies closely mirrors the altered amino acid composition observed in flies after ePR (Extended Data Fig. 4c,o). These data suggested that the reduction in storage protein Lsp2 is responsible for the shift in amino acid composition by ePR.

**Fig. 5:**
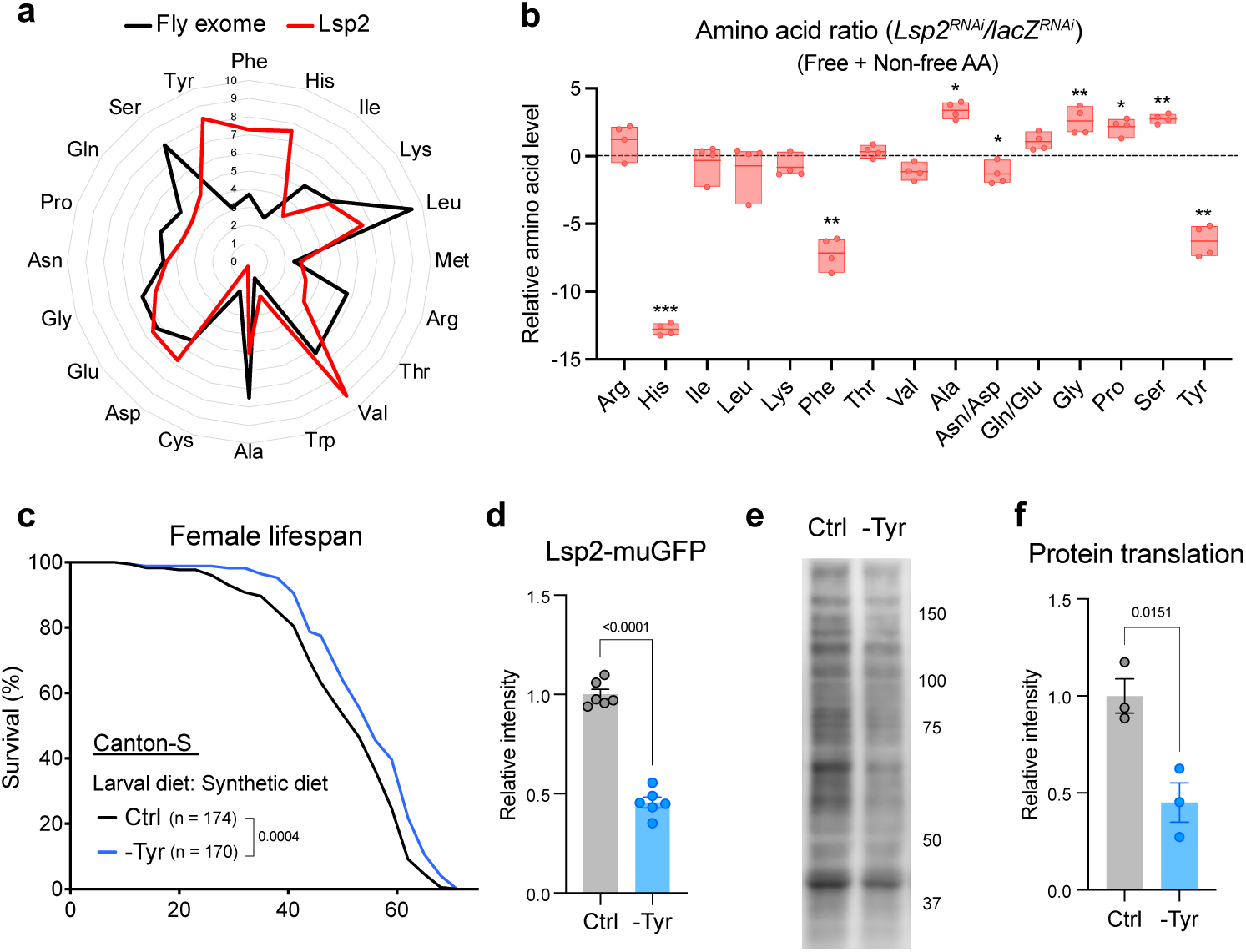
Lsp2 protein is enriched with aromatic amino acids and larval tyrosine restriction mimics ePR. **a**, Amino acid composition of Lsp2 protein and fly exome. **b**, Amino acid composition of *Lsp2*-knocked down flies or control flies (*lacZ-RNAi*) at day2 using fat body driver *r^4^-Gal4*. n = 4. **c**, Lifespan of Canton-S female flies with early tyrosine restriction. Sample sizes (n) are shown in the figure. **d**, Quantification of the *Lsp2-muGFP* reporter wandering larvae with early tyrosine restriction. n = 10. **e**,**f**, A representative image of protein translation analysis (**e**) and its quantification (**f**) in the heads of Canton-S female flies with early tyrosine restriction at day2-3. Anti-puromycin antibody is used. n = 3. For the statistics, a two-tailed Student’s *t* test (**b**,**f**), a log-rank test (**c**), or two-way ANOVA with Šídák’s multiple comparison test (**d**) was used. For the graphs, the minimum, the lower quartile, the median, the upper quartile, and the maximum points (**b**) or the mean and SEM (**d**,**f**) are shown. Data points indicate biological replicates.

Serendipitously, we have previously found that *Drosophila* is highly sensitive to Tyr intake^22,27^. Indeed, when we fed larvae with a synthetic diet specifically lacking Tyr, adult flies were long-lived (Fig. 5c). Consistently, Lsp2 expression and protein translation in early adulthood was suppressed by larval Tyr restriction (Fig. 5d-f). Taken together, these data suggest that the amount of dietary Tyr intake during larval stage regulates the amount of Tyr-enriched storage proteins, and thus it regulates adult translational capacity and lifespan.

## Discussion

In this study, we investigated how early-life dietary restriction regulates physiology in adult flies and extends lifespan. By integrating *in vivo* pSILAC, proteome analysis and transcriptome analysis, we identified the mechanism by which ePR influences early adult protein synthesis and lifespan via Lsp2. Although ePR reduces body growth, fecundity, ovariole number and insulin signalling activity, these phenotypic outcomes are not mediated by Lsp2 and thus independent from the regulation of lifespan. This fact means that the ePR longevity is not related to the reproduction-lifespan trade-off, which is consistent with the fact that both ePR and Lsp2-RNAi extends lifespan also in males.

Arylphorins, the protein group to which Lsp2 belongs, are storage proteins characteristic to holometabolous insects, which faces nutrient restriction during non-feeding pupal development^28,29^. Classic studies primarily focused on their dynamics in abundance and localisation during development. Only recently has the detailed functional study of Lsps been done, revealing its specific requirement for late-developed adult tissues^25^. It has long been believed that these storage proteins are just amino acid storage required for development, however, it is known that Lsp2 is expressed in adult flies^30^. Despite this, the role of Lsp2 in adulthood, as well as a mediator of larva-to-adult transition, has remained largely unexplored, although it was suggested that some storage proteins may serve as a reservoir for adult fecundity in some insects^31^. In our study, we show that substantial amounts of Lsp2 protein persist into adulthood. These findings indicate that Lsp2 functions not only as a developmental storage protein but also plays a pivotal role for adult physiology, through the regulation of translation capacity. Of note, this is potentially related to the fact that flies are resistant to dietary Phe deprivation as Lsp2 is rich in Phe and Tyr^32^.

It was unexpected that larval amino acids are preferentially enriched in some proteins, including ribosomal proteins. This may reflect the fact that ribosomal proteins are abundantly present and some of which are long-lived during the larval-to-adult transition. It is also possible that early adult ribosomal proteins are made of amino acids derived from storage protein. One of the storage proteins, Fat Body Protein 1 (Fbp1), is required for Lsp proteins re-uptake into the fat body to produce and utilise amino acids^25^. While the precise mechanism of Lsp2 recycling remains speculative, Lsp2 may be degraded into free amino acids through endosomal-lysosomal fusion. If this is the case, Lsp2-derived amino acids in lysosome are not equally used for protein synthesis in general. We propose that there is a “recyclome”, the group of proteins synthesised by recycled amino acids rather than newly obtained amino acids which comes directly in cytosol through transmembrane transporters. It might be associated with the fact that the lysosome serves as a key site for amino acid sensing via mTORC1. As such, Lsp2 should influence translation not only by supplying amino acid building blocks but also by modulating nutrient signalling pathways. The translation of ribosomal proteins, which contain 5’ Terminal Oligo Pyrimidine (TOP) motifs, is tightly regulated by nutrient-sensing mechanisms such as mTORC1^33^. Investigating how Lsp2 contributes to these regulatory networks could provide deeper insights into its role in translation and nutritional sensing mechanisms.

In addition to the cytosolic ribosomal proteins, *in vivo* pSILAC analysis delineates mitochondrial ribosomal proteins and mitochondrial metabolic enzymes, which is affected by ePR and *Lsp2*-RNAi. It is intriguing to test that intact mitochondria *per se* can be passed onto adult stage by escaping histolysis during pupal stage.

Mammalian species, including humans, do not have a direct ortholog of Lsp2; however, in terms of function, serum albumin and globulin can serve a similar role. For instance, albumin and Lsps share key characteristics, including detoxification through binding to low-solubility compounds, osmolality regulation, nutritional regulation, and high synthesis rates and turnover within the liver^34^. Serum albumin level is correlated with mortality risk and it can be decreased by cancer, and the albumin synthesis rate in the liver is sensitive to protein restriction^34^. Decrease in albumin leads to decrease in amino acid supply for protein synthesis in the liver and other organs. It is intriguing to consider whether albumin or globulin production is persistently affected by early-life diets. Investigating these functional parallels at the phenotypic level could provide valuable insights into conserved mechanisms of nutritional adaptation across species.

Human epidemiological studies have shown that children exposed to inadequate nutrition during development are at a significantly higher risk of developing diseases such as obesity, type II diabetes, cardiovascular disease, and schizophrenia in adulthood^35,36^. This phenomenon, termed the “Developmental Origins of Health and Disease” (DOHaD), has been extensively investigated through animal experiments, primarily in rodents. For instance, rats born to mothers fed a low-protein diet during pregnancy exhibit metabolic abnormalities, including higher insulin resistance^37^. Similarly, reducing the protein content in the maternal diet in mice during gestation has been shown to shorten the lifespan of their offspring^8^. Interestingly, mice exposed to low-protein diets during the lactation period exhibits an extended lifespan^8^. These findings suggest that the protein environment during critical developmental windows leaves a lasting imprint on the organism, inducing long-term physiological changes— both beneficial and detrimental—that persist into adulthood. Proposed mechanisms underlying this phenomenon include irreversible structural modifications in tissues, epigenetic regulation of gene expression, and the potential involvement of gut microbiota, although these mechanisms have not been fully validated^38^. Beyond these, we propose that storage proteins can encode the nutritional memory, linking developmental nutrient availability to adult physiological outcomes.

## Acknowledgements

We would like to acknowledge Kyoto Stock Center, National Institute of Genetics, Vienna Drosophila Resource Center, and Bloomington Drosophila Stock Center for reagents. We thank Valzania Luca and Pierre Leopold for Lsp2 antibody, Sebastian Sorge and Alex Gould for providing a recipe for HolFast, RIKEN BDR Genome Research Analysis Section (GRAS) for supporting RNAseq analysis, Ribeiro Carlos, Chisako Sakuma, Ayano Oi, Yusuke Kato, Yuka Fujita for technical assistance and critical reading of the manuscript. This work was supported by Japan Agency for Medical Research and Development to F.O. under Grant Number 20gm6310011, Japan Society for the Promotion of Science to F.O. under Grant Number 19H03367 and 22H02769 and to H.K. under Grant Number 22K20731, and to K.I. under Grant Number 23H04924, Japan Science and Technology Agency to F.O. under Grant Number JPMJFR2337, H.K. under Grant Number JPMJAX2226, and K.I under Grant Number JPMJFR214L.

## Author Contributions

H.K. and F.O. conceived the project. H.K. performed most of the experiments and analysed the data. K.I., C.B. and J.S. performed *in vivo* pSILAC and proteome analysis. R.O. supported the experiments. N.D. performed total amino acid quantification. H.K. and F.O. wrote the initial manuscript. F.O. supervised the study. All authors edited and approved the final manuscript.

## Materials & Correspondence

All the materials generated in this study are available upon request to F.O.

## Methods

### *Drosophila* stocks and husbandry

Stock flies and adult flies were reared on a standard yeast-based diet containing 4.5% cornmeal (Nippn Corporation), 6% brewer’s yeast (Asahi Breweries, HB-P02), 6% glucose (Nihon Shokuhin Kako), and 0.8% agar (Ina Food Industries, S-6) with 0.4% propionic acid (FUJIFILM Wako Pure Chemical Corporation, 163-04726) and 0.15% butyl p-hydroxybenzoate (FUJIFILM Wako Pure Chemical Corporation, 028-03685). Flies were maintained at 25°C. To allow synchronised development and constant density, embryos were collected using agar plates (2.3% agar, 1% sucrose, and 0.35% acetic acid) with a live yeast paste and 10–15 μL of the embryos were spread onto bottles.

Fly lines used in this study were Canton-S, *w^Canton-S^*(*w^iso^*^31^ eight times backcrossed with Canton-S), *4E-BP^intron^-dsRed*^23^, *Cg-Gal4* (Bloomington Drosophila Stock Center (BDSC), 7011, eight times backcrossed with *w^Canton-S^*), *tub-Gal80^ts^* (BDSC 7017), *r*^4^*-Gal4* (Bloomington 33832, eight times backcrossed with *w^Canton-S^*), *fit-Gal4*^20^, UAS-2×EGFP (BDSC 6874), *UAS-lacZ-RNAi* (from Dr. Carthew R., eight times backcrossed with *w^Canton-S^*)*, UAS-Lsp2-RNAi* (National Institute of Genetics (NIG), 6806R-2, eight times backcrossed with *w^Canton-S^*), *UAS-Lsp1α-RNAi* (Vienna Drosophila Resource Center (VDRC), 14898), *UAS-Fbp2-RNAi* (VDRC, 33173), *Lsp2-muGFP* (created by CRISPR/Cas9 system, shown as below).

To generate *Lsp2-muGFP* flies, we used the CRISPR/Cas9 system to insert *monomeric ultrastable GFP* (*muGFP*) at the C-terminus of the *Lsp2* gene^39^. The codons of *muGFP* were optimised for expression in *Drosophila melanogaster*. These modifications and insertions into the EcoRI/XbaI site in the pUC57 vector were performed by GenScript. An sgRNA target site of *Lsp2* was selected using CRISPR Optimal Target Finder^40^. Complementary oligonucleotides with overhangs were annealed and cloned into the BbsI-digested U6b vector using a DNA ligation kit (Takara, 6023).

Sense strand: TTCGGTCCAGGATCTAGACCACAT

Antisense strand: AAACATGTGGTCTAGATCCTGGAC

The targeting vector was constructed by inserting linker sequence (GGTGGATCTGGAGGTTCCGGCGGCTCAGGGGGTAGT) and *muGFP* between 500 bp of *Lsp2* homology arms with a silence mutation in the PAM sequence next to the sgRNA site (from TGG to TTG). First, linker sequence with *muGFP* and homology arms were PCR-amplified using Q5 High-Fidelity 2× Master Mix (New England BioLabs, M0492L). Primers for PCR were designed using the NEBuilder Assembly Tool. The gel-purified PCR products were cloned into the EcoRI-digested pBluescript II SK(+) vector using NEBuilder HiFi DNA Assembly Master Mix (New England BioLabs, E2621X). The mixture of pU6b-sgRNA and targeting vector was microinjected into *w*^1118^*; attP40{nos-Cas9}/CyO* embryos by WELLGENETICS. The F0 adults were crossed with balancer lines, and the correct *muGFP*-inserted lines were selected by PCR amplification and sequencing of the target locus. The primers used for PCR are listed in Table S1.

### Dietary manipulations

For larval protein restriction, baker’s yeast (Lesaffre, Saf-instant, red), 6% glucose (FUJIFILM Wako Pure Chemical Corporation, 049-31165), 1% agar (FUJIFILM Wako Pure Chemical Corporation, 010-15815), 0.3% propionic acid (FUJIFILM Wako Pure Chemical Corporation, 163-04726), and 0.15% methyl p-Hydroxybenzoate (FUJIFILM Wako Pure Chemical Corporation, 132-02635) were used to produce 8%, 2%, or 1% yeast diet. Embryos were collected within four hours of egg laying in a cage (Flystuff, 59-101 or 59-100). The embryos (10–15 μL) were spread by micropipette onto a surface of 8% yeast diet and raised until late 2^nd^ instar larva. At around 66 hr after egg laying, the larvae were floated up using 30% glycerol and transferred to each diet. Normally, 150–200 adult flies were obtained per bottle. After eclosion, adult flies were collected to standard yeast-based diet.

Chemically defined diet, or holidic medium, was used for larval tyrosine restriction^41^. AA concentrations were modified according to the optimised diet through exome-matching^16^. The sugar source (not sucrose but glucose), and the amount of agar and preservatives were modified as a previous study^22^. Embryos were spread by micropipette onto a surface of standard yeast-based diet and raised until late 2^nd^ instar larvae. At around 66 hr after egg laying, the larvae were floated up using 30% glycerol and transferred to each holidic medium. After eclosion, adult flies were collected to standard yeast-based diet.

### Analysis of developmental speed, eclosion rate, and body weight

Embryos were spread onto the six bottles and the pupal number of each bottle was counted every several hours. The number of eclosed adult flies per pupal number was calculated as eclosion rate. For the body weight measurement, each single fly was anesthetised by CO_2_ and placed onto a microbalance (METTLER TOLEDO, XPR2).

### Lifespan analysis

Adult flies were allowed to mate for two days after eclosion in a bottle. Subsequently, 30 female or male flies were allocated to each vial. Six vials containing 30 flies each were prepared for each condition. The flies were maintained at 25°C under 60% humidity and a 12 h light:12 h dark cycle. Flies were transferred to new vials two to three times per week and the number of dead or censored flies were counted.

### Fecundity analysis

Similar to the procedure for lifespan analysis, at two days after eclosion, 17 female flies and 17 male flies were placed in vials containing a specific diet. Two vials containing a total of 34 flies per vial were prepared for each condition. After one week, flies were anesthetised quickly with CO_2_ and reallocated to 10 vials containing each diet with three females and three males per vial. After 24 h, the number of eggs laid on the medium was manually counted.

### Western blot analysis

The abdominal carcass from eight female flies or head from 12 female flies were dissected in PBS and homogenised in 50 μL of RIPA buffer (FUJIFILM Wako, 188-02453) supplemented with a protease inhibitor (FUJIFILM Wako, 165-26021) and phosphatase inhibitor cocktails (Roche, 4906845001). The supernatant was collected after centrifugation and the protein amount was quantified by BCA assay (FUJIFILM Wako, 164-25935). The samples were mixed with 6× SDS–PAGE sample buffer (nacalai, 09499-14) and 15–20 μg proteins were subjected to standard SDS–PAGE. Gels were transferred to the PVDF membrane and blocked by Everyblot blocking buffer (Bio-rad, 12010020). Primary antibodies used in the study were anti-α + β tubulin (Abcam, ab44928, 1:1000 dilution), anti-histone H3 (CST, 14269S, 1:1000 dilution), anti-phospho-Akt (CST, 4060T, 1:1000 dilution), anti-total Akt (CST, 9272S, 1:1000 dilution), anti-phospho-eIF2α (CST, 3398S, 1:1000 dilution), anti-total eIF2α (abcam, ab26197, 1:1000 dilution), and anti-Lsp2^25^. HRP-conjugated secondary antibodies were anti-mouse IgG, HRP-linked antibody (CST, 7076S, 1:1000 dilution), anti-rabbit IgG, HRP-linked antibody (CST, 7074S, 1:1000 dilution), and anti-rat IgG, HRP-linked antibody (Jackson ImmunoResearch, 81211, 1:1000 dilution). The signals were visualised by chemiluminescence using Immobilon (Millipore, WBLUF0100) and detected by Amersham ImageQuant 800 (Cytiva).

### Protein synthesis assay

Protein synthesis was monitored based on the reported SUnSET assay^42^. Adult flies were fed with 600 μM puromycin (FUJIFILM Wako, 160-23154) in standard diet for 24 hours. Twelve heads of female flies were collected and lysed in the RIPA buffer (FUJIFILM Wako, 188-02453) supplemented with a protease inhibitor (FUJIFILM Wako, 165-26021). 20 μg of protein samples was mixed with a 6× SDS-PAGE sample buffer and subjected to the standard SDS-PAGE technique using 10–20% gradient gel (FUJIFILM Wako, 198-15041). Gels were transferred to the PVDF membrane and blocked by 4% skimmed milk. The incorporated puromycin was visualised by the western blot analysis using anti-puromycin (Millipore, MABE343, 1:1000 dilution) and anti-mouse IgG2a (Jackson ImmunoResearch, 115-035-206, 1:15000 dilution).

### RNA sequencing analysis and quantitative RT‒PCR analysis

For RNA sequencing analysis, we dissected 12 heads and 8 abdominal carcasses from female flies. Gut, Malpighian tubules, and ovaries were carefully removed from the abdomen to prevent contamination. Total RNA was purified from the samples using a ReliaPrep RNA Tissue Miniprep kit (z6112, Promega Corporation). Four samples were prepared for each experimental group. Library preparation was performed by the RIKEN BDR Technical Support Facility using an Illumina Stranded mRNA Prep Ligation kit (96 Samples) (20040534, Illumina K.K.) and IDT^®^ for Illumina^®^ RNA UD Indexes Set C Ligation kit (20091659, Illumina K.K.). The optimal number of PCR cycles was determined by real-time PCR using KAPA SYBR FAST qPCR Master Mix (KK4603, F. Hoffmann-La Roche, Ltd.). The library quality was verified using the TapeStation HS D1000 assay. The library samples were forwarded to AZENTA for RNA sequencing using an Illumina NovaSeq 6000 (Illumina K.K.). The paired-end 150 bp sequence data were analysed as follows: a quality check of the raw reads was performed by FastQC (v0.12.1)^43^ and MultiQC (v1.24)^44^. The raw reads were then filtered to remove the first base (T), adaptors, and low-quality bases using Trim Galore (v0.6.10)^45^. Filtered reads were aligned to the *Drosophila* genome (BDGP6.46) using Hisat2 (v2.2.1) ^46^. The read counts were calculated using StringTie (v2.2.1)^47^. Differentially expressed genes were identified using edgeR (v4.0.6)^48^. RNA-sequencing data have been deposited at the DDBJ under accession number DRR622306-DRR622313, DRR626770-DRR626793.

For quantitative RT‒PCR analysis, total RNA was purified from 6 heads or 8 abdominal carcasses of female flies, or 4 whole bodies of larvae as described above using a ReliaPrep RNA Tissue Miniprep kit (z6112, Promega Corporation). The cDNA was synthesised from 500 ng of DNase-treated total RNA using Revertra Ace Master mix (FSQ-201, Toyobo Co., Ltd.). Quantitative RT‒PCR was performed using Taq Pro Universal SYBER qPCR Master Mix (Q712-02-AA, Vazyme Biotech Co., Ltd.) and qTOWER^3^ G (Analytik Jena GmbH+Co. KG). The ΔΔCt method was used, with *RNA pol2* serving as the internal control. Primer sequences are listed in Table S1.

### Imaging analysis

To analyse ovary size, dissected ovaries were placed in phosphate buffer saline (PBS) on a silicone pad. Images were captured using a fluorescence stereomicroscope (MZ10F, Leica Microsystems GmbH). The area of the ovary from the top view was measured using Fiji software^49^. Ovariole number was counted by pulling out the each ovariole from ovaries by forceps and averaging the left and right ovariole numbers. For whole body reporter fluorescence, flies were immobilised on a CO_2_ pad and the red fluorescent protein (RFP) or GFP fluorescence images were captured using a fluorescence stereomicroscope (MZ10F, Leica Microsystems GmbH). To analyse ATF4 reporter fluorescence in the abdomen, flies were imaged from the lateral side to minimise the interference of the basal fluorescence in the gut and Malpighian tubules. A region between stripes of the dorsal abdomen was selected as region of interest (ROI) and the fluorescence was quantified using Fiji software^49^. The expression level of the *fit>GFP* reporter was quantified by measuring the fluorescence of the entire abdomen, delineated by elliptical selections using the Fiji software package^49^. The expression level of the *Lsp2-muGFP* reporter was quantified by measuring the fluorescence of whole bodies of larvae, pupae, and adults.

### Total amino acid quantification

Four whole flies were collected in each crimp glass vial (Crimp Top Vials, P/N: 03-CVG, Chromacol, UK) after body weight measurement was conducted using a microbalance (METTLER TOLEDO, XPR2). Four crimp glass vial samples were prepared for each condition and stored at −80°C. The amino acid amount was normalised by the body weight of the flies. After vacuum drying, each crimp vial was placed in borosilicate glass vials (P/N: 224832, Wheaton, NJ USA). 200 μL of 6N HCl and a small phenol crystal were then added to the outside of the crimp vials. For Trp analysis, 4N methanesulfonic acid with 0.2% (w/v) tryptamine (FUJIFILM Wako Pure Chemical Co., Japan) was added in the crimp vials instead of the HCl solution. After evacuating for a few minutes, the vial was sealed with a Mininert valve (No. SC-24, P/N: 10130, Pierce, IL USA) and heated in a heating bath at 110°C for 20 hours. All procedures for amino acid analysis using precolumn derivatisation with 6-aminoquinolyl-N-hydroxysuccinimidyl carbamate were as described in^50^.

### In vivo pulse SILAC proteome analysis

Standard proteome analysis was performed on a total of 12 heads of day 2 female flies. These flies were homogenised in 50 μL of RIPA buffer (FUJIFILM Wako, 188-02453) supplemented with a protease inhibitor (FUJIFILM Wako, 165-26021). In the isotope labelling proteome experiment, larvae were fed a complete HolFast diet^51^ with lysine and arginine substituted with heavy isotope-labelled lysine (+8, Cambridge Isotope Laboratories, Inc (CIL), CNLM-291-H-0.1) and arginine (+10, CIL, CNLM-539-H-0.1). These amino acids are recognised and cleaved by trypsin, resulting in peptides that terminate with a single lysine or arginine residue at their C-terminus. Subsequent to eclosion, adult flies were transferred to a standard yeast diet at day 0. Two days or 12 days after eclosion, the flies were transferred to holidic medium^41^ (exome matched version^16^) with lysine and arginine substituted with medium isotope-labelled lysine (+4, Cambridge Isotope Laboratories, Inc (CIL), DLM-2640-0.1) and arginine (+6, CIL, CLM-2265-H-0.1). Following a four-day feeding period with isotope-labelled amino acids, 12 heads of female flies were homogenised in 50 μL of RIPA buffer (FUJIFILM Wako, 188-02453) supplemented with a protease inhibitor (FUJIFILM Wako, 165-26021). Six samples were prepared for standard proteome analysis, while three samples were prepared for isotope-labelled proteome analysis.

For the proteomics analysis, protein extracts in RIPA buffer were sonicated using a BioRuptor with 10 cycles on high power with 60 sec on followed by 30 sec off. Then the samples were centrifuged at 18,000 x g for 20 min at 4°C and 10 μL of the sample supernatants were taken. Detergent concentration in the samples was decreased using methanol-chloroform extraction^52^. Then 105 µL 50 mM ammonium bicarbonate buffer was added to the protein pellets. Protein amounts were quantified using 5 µL with a BCA assay kit (Thermo Fisher). The average protein amount in 100 µL was 12 µg. The cysteine disulfide bonds were reduced with 10 mM Tris (2-carboxyethyl) phosphine Hydrochloride (TCEP-HCl) at 37°C for 30 min, and then the cysteines were alkylated with 50 mM 2-Chloroacetamide (CAA) at RT for 30 min. For protein digestion, 300 ng Lys-C was added, and the samples were incubated at 37°C for one hour. Then 300 ng trypsin was added, and the samples were incubated at 37°C for 16 h. The next day, samples were acidified with ∼0.5% TFA (final concentration). Tryptic peptides were desalted with polystyrene-divinylbenzene, reversed-phase sulfonate (SDB-RPS) StageTips^53^.

For sample measurement, peptides were eluted from SDB-RPS StageTips with 40 µL 5% ammonia, 15% water and 80% acetonitrile, and vacuum dried. Afterwards, the peptides for each sample were dissolved in 10 µL 0.1% formic acid, 3% acetonitrile, 97% water. A nanoLC/MS/MS system comprising a Vanquish Neo UHPLC (Thermo Fisher Scientific) and an Orbitrap Eclipse mass spectrometer (Thermo Fisher Scientific) was employed. The mobile phases consisted of (A) 0.1 % formic acid and (B) 0.1 % formic acid and 80 % acetonitrile. Peptides were loaded on a self-made 20 cm fused-silica emitter (75 µm inner diameter) packed with ReproSil-Pur C18-AQ (1.9 µm, Dr. Maisch) and separated by a linear gradient (2−40% B in 90 min, 40−99% B in 5 min, and 99% B for 10 min) at the flow rate of 300 nL/min. All MS1 spectra were acquired over 375–1500 *m/z* in the Orbitrap analyzer (resolution = 60,000, maximum injection time = 50 msec, and automatic gain control = 4e5). For the subsequent MS/MS analysis, precursor ions were selected and isolated in a top-speed mode (cycle time = 3 sec and isolation window = 1.6 *m/z*), activated by higher-energy collisional dissociation (normalized collision energy = 28), and detected in the Orbitrap analyzer (resolution = 30,000, maximum injection time = 54 msec, and auto gain control = 5e4). Dynamic exclusion time was set to 30 sec.

Raw data files were analysed and processed by MaxQuant^54^ (v2.1.0.0), and the database search was performed with Andromeda^55^ against the database of 22,034 *Drosophila melanogaster* (UP000000803) Swiss-Prot reviewed and TrEMBL unreviewed canonical and isoform proteins downloaded from UniProt on September 30, 2024, with common contaminants and enzyme sequences. Search parameters included two missed cleavage sites and variable modifications such as L-(13C6,15N4)-arginine (Arg10 = heavy), L-(13C6,15N2)-lysine (Lys8 = heavy), L-(13C6)-arginine (Arg 6 = medium), L-(D4)-lysine (Lys 4 = medium), methionine oxidation, and protein N-terminal acetylation. Cysteine carbamidomethylation was set as a fixed modification. The peptide mass tolerance was 4.5 ppm, and the MS/MS tolerance was 20 ppm. The false discovery rate was set to 1% at the peptide spectrum match evel and protein level. For protein-level quantification, ‘unique + razor’ peptides were used. For the SILAC-based protein quantification, a minimum of one ratio count (unique peptide ion) was used for quantification, and the ‘re-quantify’ and ‘match between runs’ functions were employed. Raw H/M ratios were used for quantification. To estimate abundance of light-, medium-, and heavy-labelled proteins within the samples, intensity-based absolute quantification (iBAQ) which computes the sum of all the peptides intensities divided by the number of theoretically observable peptides was used^56^. Top 25% abundant heavy-labelled proteins were selected and used for GO analysis.

### Proteome analysis

For the proteomics analysis, protein extracts in RIPA buffer were sonicated using a Bioruptor BR-11 on high power for 10 cycles with one min on followed by 30 sec off. The samples were then centrifuged at 10,000 x g for 10 min at 4°C. The Phase Transfer Surfactant (PTS) method of sample preparation was followed^57,58^, 10.6 µL supernatant was taken from each sample and mixed with 144.4 µL PTS buffer (12 mM SDC, 12 mM SLS, 100 mM Tris.Cl pH 8.5). A 5 µL aliquot of each sample was taken for quantification using a BCA assay. The average protein amount was about 35 µg in 150 µL protein extract. TCEP was added at 10 mM and the samples were incubated at 37°C for 30 min, then CAA was added at 50 mM, and the samples were incubated at 25°C for 30 min. Then 600 µL 50 mM ammonium bicarbonate was added, followed by 700 ng Lys-C and 700 ng trypsin for protein digestion at 37°C for 16 h. The following day, 780.3 µL ethyl acetate to each sample followed by addition of TFA to 0.75% final concentration. The samples were vortexed for 2 min then centrifuged at 12,000 x g, for 5 min at RT. The supernatant was discarded. The samples were desalted with SDB-RPS StageTips.

The samples were eluted from the SDB-RPS StageTips, vacuum dried, and then dissolved in around 17.5 µL 0.1% formic acid, 3% acetonitrile, 97% water. A Q Exactive Plus mass spectrometer, together with an EASY-nLC 1200, and NanoSpray Flex Ion Source (Thermo Fisher, USA) were used for sample measurement. Analytical columns were 75-µm I.D., and contained 1.9-µm particle C18 particles, with 18-cm filling length. The gradient conditions were as follows with the percentage of acetonitrile as indicated: 0–1 min, 0.0-4.0%; 1–110 min, 4.0–32.0%; 110–112 min, 32.0–76.0%; 112–120 min, 76.0%; 120–121 min, 76.0-4.0%; 121-136 min, 4.0%. The flow rate was 150 nL/min. The ion transfer capillary temperature was 250°C and the spray voltage was 2.0 kV. MS acquisition conditions were as previously described. Briefly, a full-MS scan was collected from 385 to 1015 m/z at 35,000 resolution, an AGC target of 1e6, with a maximum IT of 55 ms. This was followed by 76 DIA scans at 17,500 resolution, an AGC target of 1e6, with a maximum IT of 55 ms, a default charge state of 3, a loop count of 38 and isolation windows of 16.0 m/z, a fixed first mass of 150.0 m/z and 27% HCD collision energy. Two staggered series of 38 windows were used 408.4355 to 1000.7047 then 400.4319 to 992.70110. All spectrum data were centroid type. Raw data files acquired with data-independent acquisition were analysed with DIA-NN software^59^, version 1.9.2. A predicted spectral library was first made using DIA-NN, which was annotated with two sequence databases: a database of 22,034 *Drosophila melanogaster* (UP000000803) Swiss-Prot reviewed and TrEMBL unreviewed canonical and isoform proteins downloaded from UniProt on September 30, 2024, and a database of common mass spectrometry contaminant proteins, which was included with the DIA-NN software package. For the data analysis, output was filtered at 0.01 FDR, N-terminal methionine excision enabled, the maximum number of missed cleavages was set to 1, cysteine carbamidomethylation was enabled as a fixed modification, protein inference for generating a subset of all protein IDs matched to the precursor was set to relaxed mode, and the empirical library generation mode was set to identifications, retention time and ion mobility profiling. The output from the data analysis included a protein group matrix file of the relative protein abundancies.

### Statistical analysis

Statistical analysis was performed using GraphPad Prism 10 (MDF Co., Ltd.). The sample size was determined empirically. To eliminate biological bias, the flies were randomly distributed onto each diet. All data points were biological, not technical, replicates. No data were excluded. Since the experiment planner (HK) and the experimenter were the same person, most experiments were not conducted in a blinded manner. For the experiments on lifespan, a different person (RO) conducted the experiments without prior bias. An unpaired and two-sided Student’s *t* test was used to compare samples. One-way ANOVA with Holm-Šídák’s multiple comparison test was used to compare groups. One-way ANOVA with Dunnett’s multiple comparison test was used to compare against a control sample. All experimental results were repeated at least twice to confirm reproducibility. Bar graphs are drawn as the mean and standard error of the mean (SEM). For the proteome analyses of ePR and Lsp2 RNAi experiments, we used only proteins that were quantified in all of the 6 biological replicates of at least one condition for further analyses. Quality check of the proteomic data was done by an R package, DEP^60^, and shown in Extended Data Fig.5, 11. Missing values were imputed using random draws from a Gaussian distribution centred around a minimal intensity value. Two-sided Wilcoxon rank-sum test was used to compare groups (all vs cytosolic ribosomal proteins or mitochondrial ribosomal proteins).

## Data availability

All relevant data are available from the authors upon request. The NGS data are available under accession numbers DRR622306-DRR622313, DRR626770-DRR626793. The proteomics data have been deposited to the ProteomeXchange Consortium via the jPOST^61^ partner repository with the dataset identifier PXD062930 (JPST003763).

## Code availability

All analysis codes are available from authors upon request.

---

**Extended Data Fig. 1:**
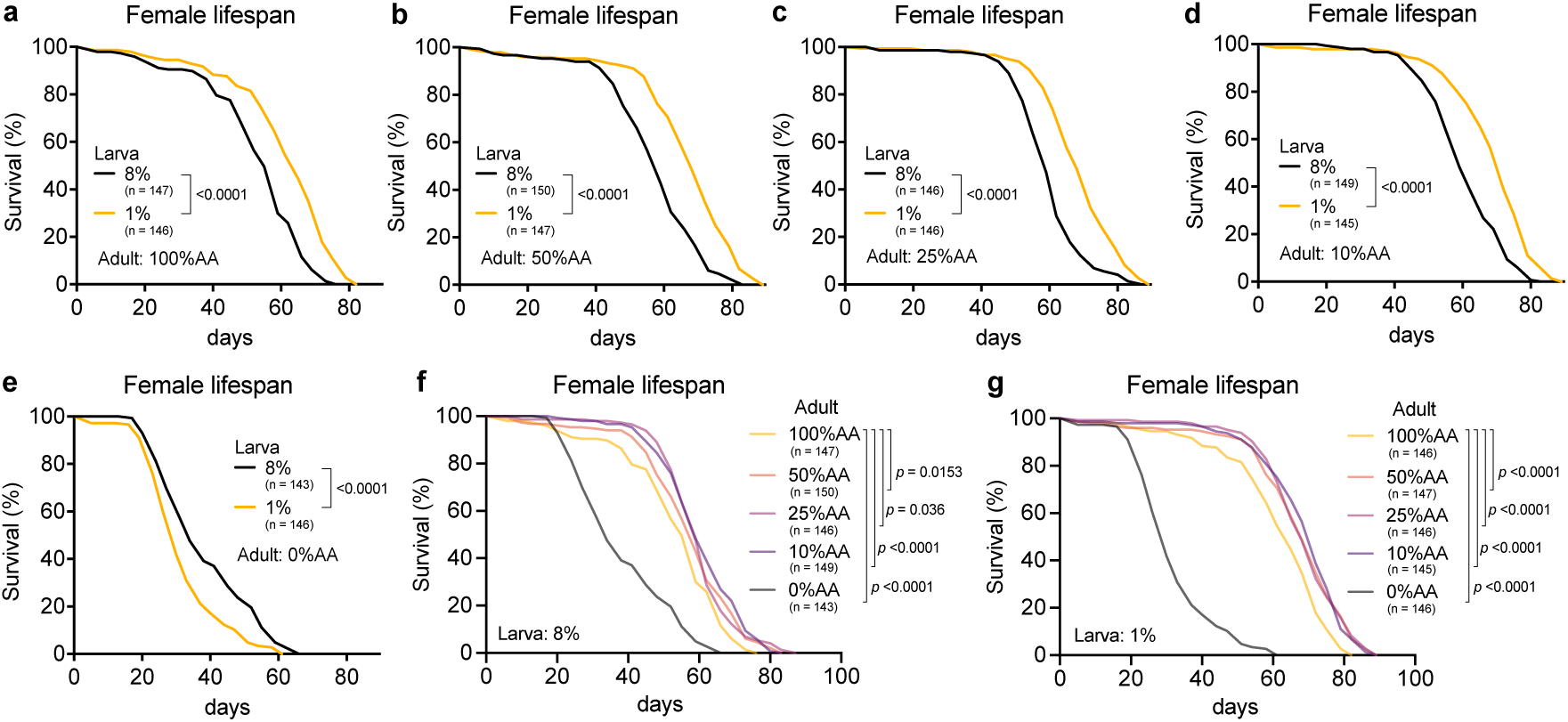
Lifespan of female flies under different adult diet with or without ePR. **a**-**e**, Lifespan of Canton-S female flies with ePR and adult feeding of synthetic diet varying the amount of amino acid (100% (**a**), 50%(**b**), 25% (**c**), 10% (**d**), and 0% (**e**)). **f**,**g**, The comparison of lifespan of **a**-**e** between adult amino acid concentration with control diet (**f**) or with ePR (**g**). Sample sizes (n) are shown in the figure. For the statistics, a log-rank test was used.

**Extended Data Fig. 2:**
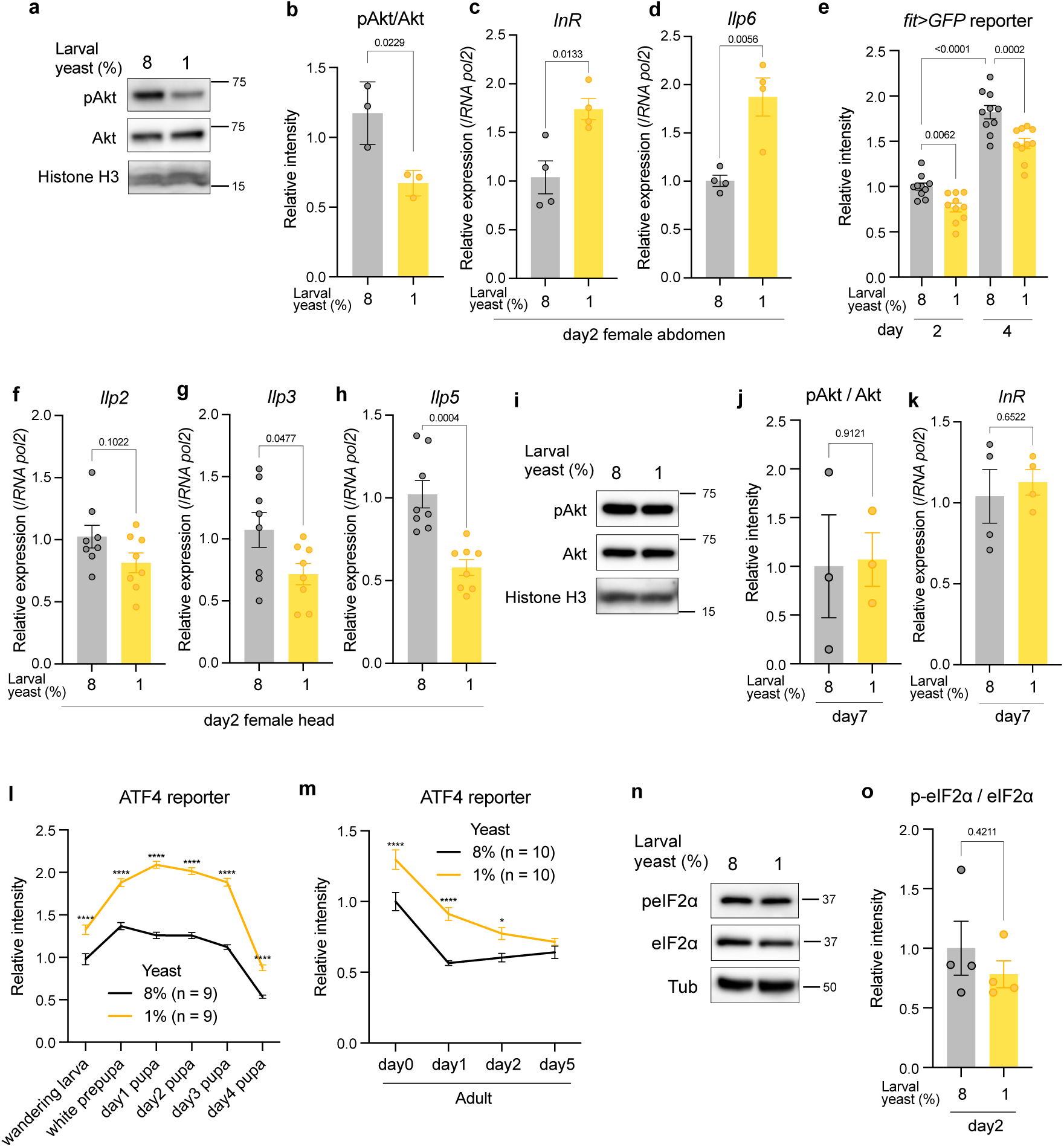
Nutritional signalling of the adult flies after ePR. **a**,**b**, Representative images of western blot analysis (**a**) and its quantification (**b**) of adult abdominal carcasses of Canton-S female flies with ePR at day2. Antibodies used are anti-phosphorylated Akt (pAkt), anti-Akt, and anti-histone H3 (loading control). n = 3. **c**,**d**, Quantitative RT‒PCR analysis of *InR* (**c**) and *Ilp6* (**d**) in the abdominal carcass of Canton-S female flies with ePR at day2. n = 4. **e**, Quantification of *fit>GFP* reporter in the abdomen of female flies with ePR at day2 and day4. n = 10. **f**-**h**, Quantitative RT‒PCR analysis of *Ilp2* (**f**), *Ilp3* (**g**), and *Ilp5* (**h**) in the heads of Canton-S female flies with ePR at day2. n = 8. **i**,**j**, Representative images of western blot analysis (**i**) and its quantification (**j**) of adult abdominal carcasses of Canton-S female flies with ePR at day7. Antibodies used are anti-phosphorylated Akt (pAkt), anti-Akt, and anti-histone H3 (loading control). n = 3. **k**, Quantitative RT‒PCR analysis of *InR* in the heads of Canton-S female flies with ePR at day7. n = 4. **l**,**m**, Quantification of ATF4 reporter (*4E-BP^intron^-dsRed*) fluorescence of larvae and pupae (**l**) and female adults (**m**) with ePR. n = 9 (**l**) or 10 (**m**). **n**,**o**, Representative images of western blot analysis (**n**) and its quantification (**o**) of adult abdominal carcasses of Canton-S female flies with ePR at day2. Antibodies used are anti-phosphorylated eIF2α (peIF2α), anti-eIF2α, and anti-tubulin α+β (loading control). n = 4. For the statistics, a two-tailed Student’s *t* test (**b**-**d**,**f**-**h**,**j**,**k,o**), one-way ANOVA with Holm-Šídák’s multiple comparison test (**e**), or two-way ANOVA with Šídák’s multiple comparison test (**l**,**m**) was used. \**P* < 0.05; \*\*\*\**P* < 0.0001. For all graphs, the mean and SEM are shown. Data points indicate biological replicates.

**Extended Data Fig. 3:**
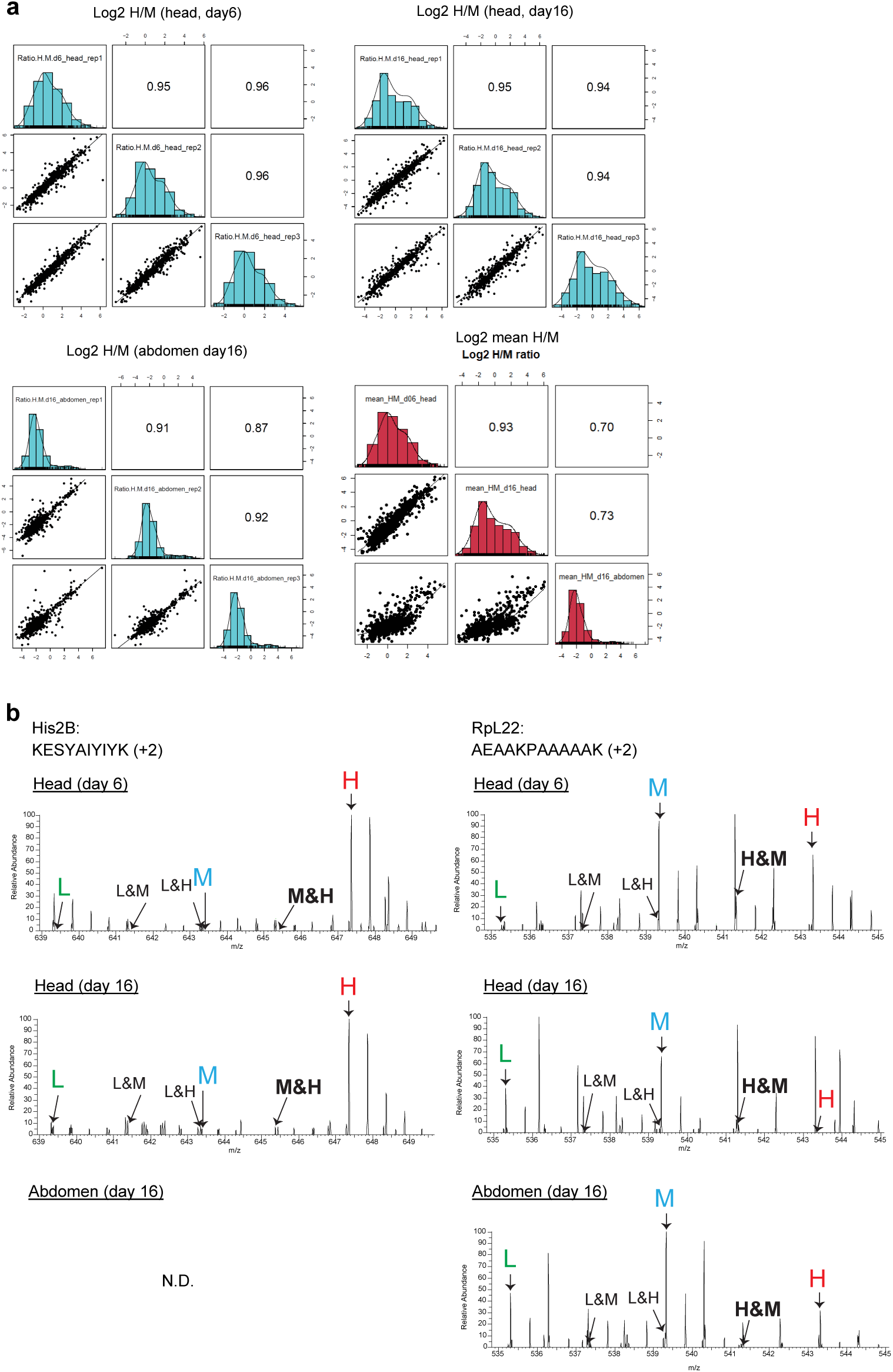
Quality control of proteome analysis and chimera protein analysis. **a**, Multi-scatter plots showing correlation of log2 H/M ratios between biological replicates, indicating high-reproducible datasets. **b**, MS1 spectra of missed-cleaved tryptic peptides which contain two labelled lysine derived from a slow (His2B) and intermediate (RpL22) turnover protein. Due to amino acid recycling after protein degradation, chimera peaks containing L&M, L&H, or M&H amino acids can be observed. Indeed, partially labelled chimera peaks are observed, but these peaks are small compared to the fully labelled peaks.

**Extended Data Fig. 4:**
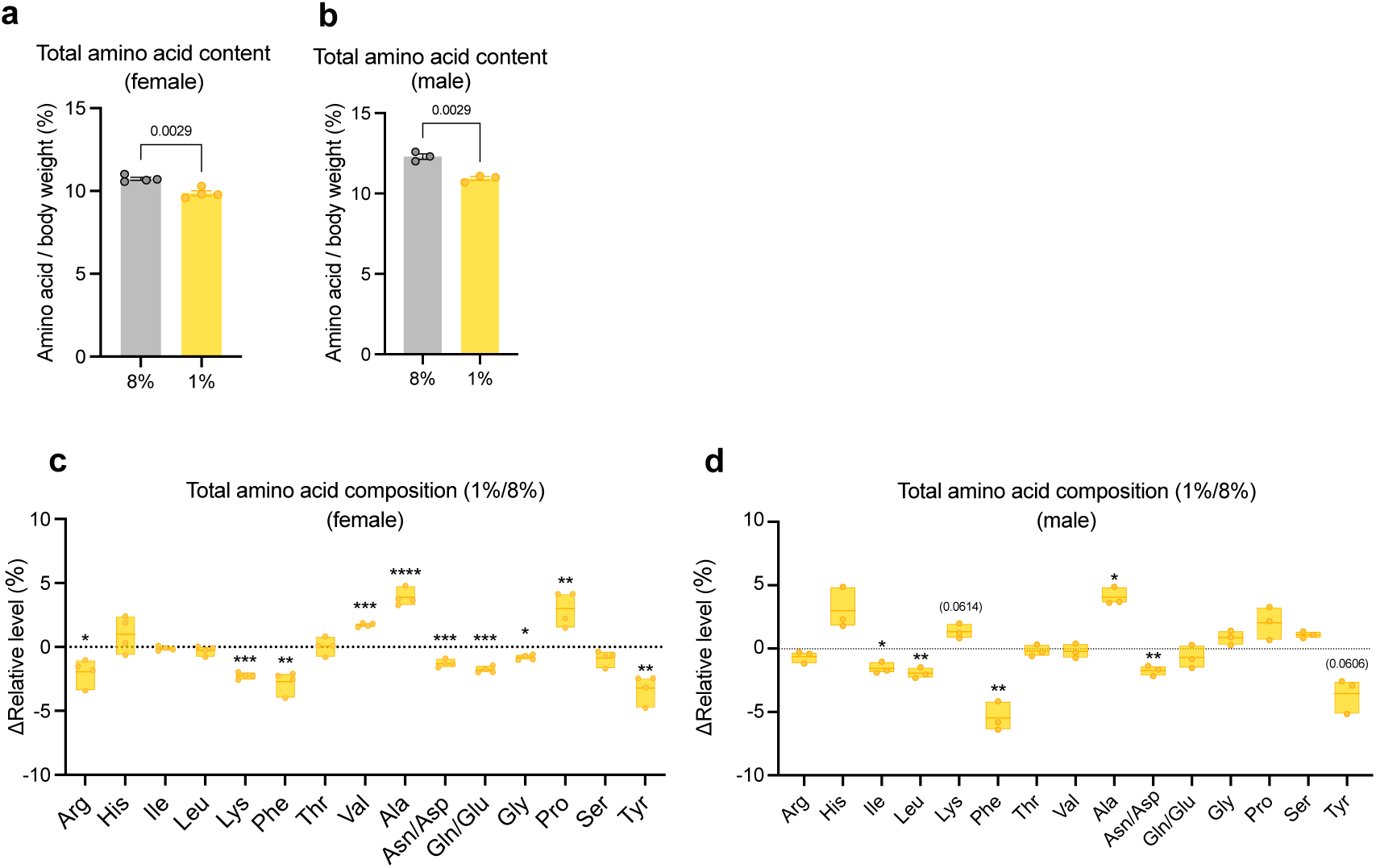
Total amino acid contents of adult flies with ePR. **a**-**d**, The total amino acid content (**a**,**b**) and composition (**c**,**d**) of Canton-S day3 female (**a**,**c**) or day0 male (**b**,**d**) flies with ePR. n = 4 (**a**,**c**) or 3 (**b**,**d**). For the statistics, a two-tailed Student’s *t* test (**a**-**d**) was used. \**P* < 0.05; \*\**P* < 0.01; \*\*\**P* < 0.001; \*\*\*\**P* < 0.0001. For the graphs, the mean and SEM (**a**,**b**) or the minimum, the lower quartile, the median, the upper quartile, and the maximum points (**c**,**d**) are shown. Data points indicate biological replicates.

**Extended Data Fig. 5:**
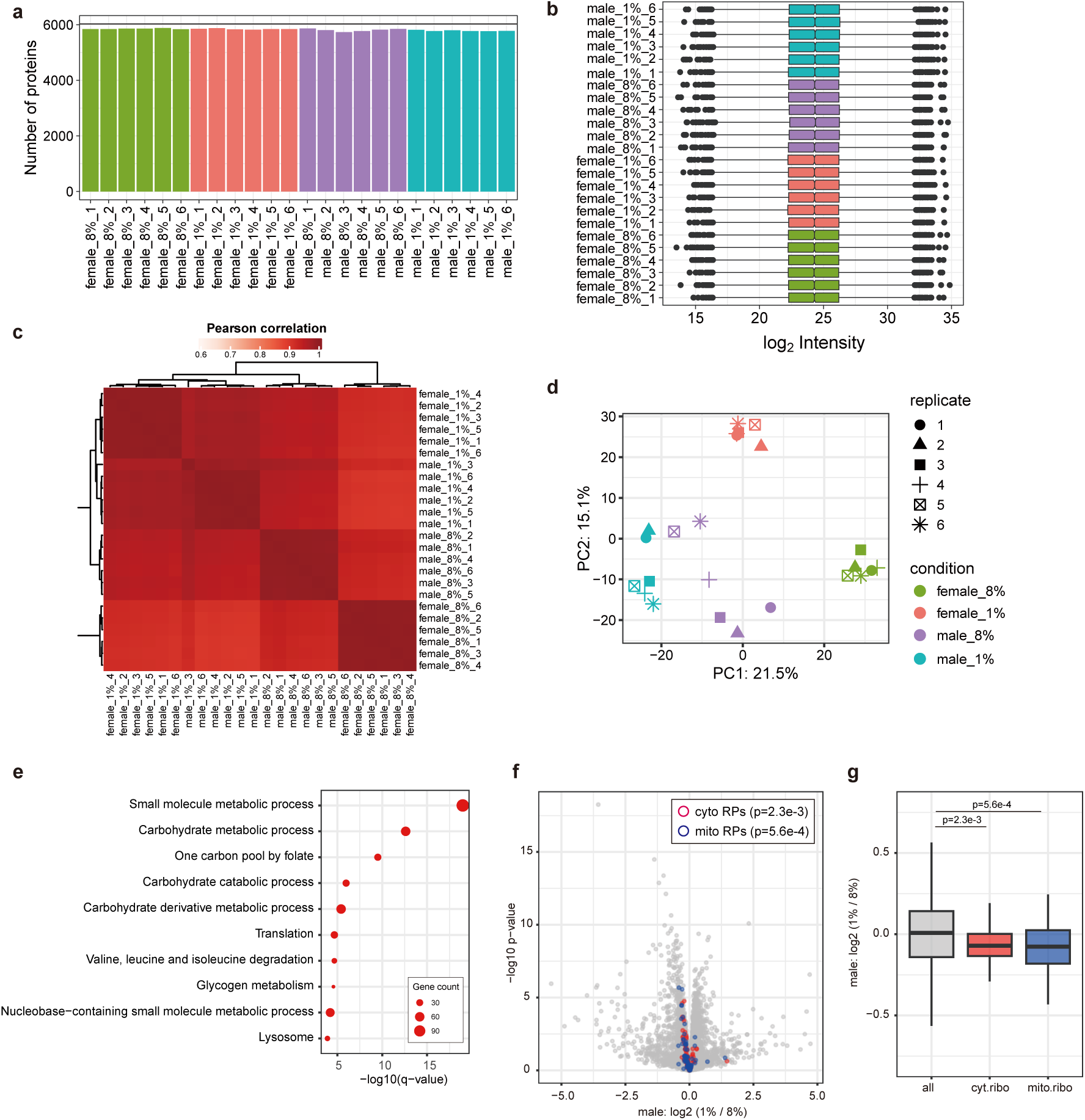
Proteome analysis of adult flies with ePR. **a**, The number of proteins quantified in each sample: Of the 6,559 proteins identified in total, 6,025 proteins that were consistently quantified across all six biological replicates in at least one condition were included in the analysis. **b**, Overview of log2 protein intensity of individual samples. **c**, Correlation matrix showing Pearson correlation of log2 protein intensity. **d**, PCA plot of individual samples. **e**, GO enrichment analysis of proteins which expression was significantly decreased by ePR in heads of Canton-S male flies at day2 (p<0.01 and log2 1% / 8% <-0.1). **f**, A volcano plot showing mean log2 fold change (1% / 8%) and −log10 p-value of proteins in the head of Canton-S male flies with ePR at day2. The cytosolic and mitochondrial ribosomal proteins are indicated by red and blue circles, respectively. **g**, A box plot showing log2 fold change (1% / 8%) of proteins grouped into three categories: all proteins, cytosolic and mitochondrial ribosomal proteins. For the statistics, two-sided Wilcoxon rank-sum test (**g**) was used. For the graph, the minimum, the lower quartile, the median, the upper quartile, and the maximum points (**g**) are shown. Data points indicate biological replicates.

**Extended Data Fig. 6:**
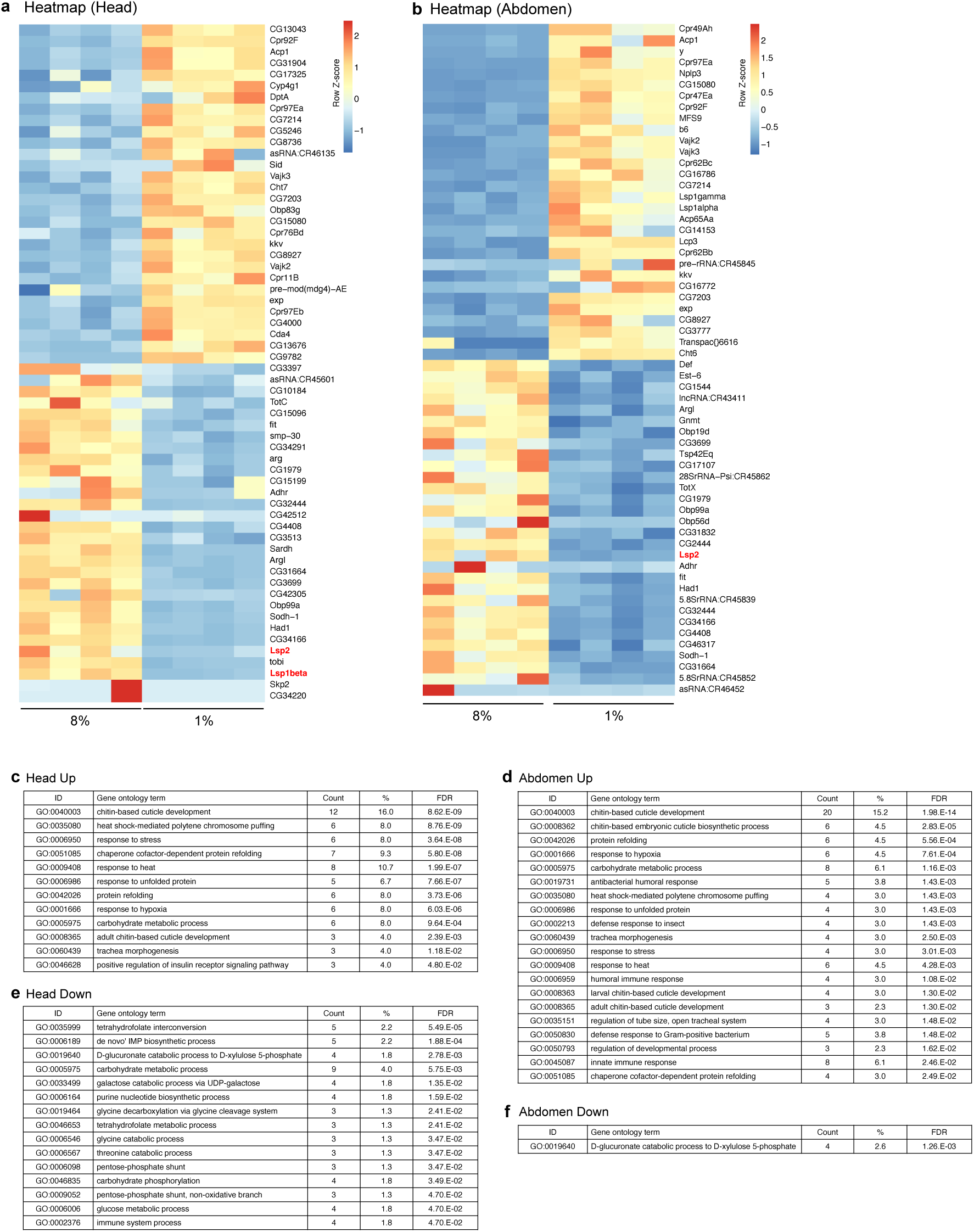
Transcriptome analysis of adult flies with ePR. **a**,**b**, Heat map of head (**a**) and abdominal carcasses (**b**) of Canton-S female flies with ePR at day2. Top 30 up-regulated and down-regulated genes are shown. **c**-**f**, Gene ontology analysis of up-(**c**,**d**) or down-(**e**,**f**) regulated genes in the head (**c**,**e**) or abdomen (**d**,**f**) of Canton-S female flies with ePR.

**Extended Data Fig. 7:**
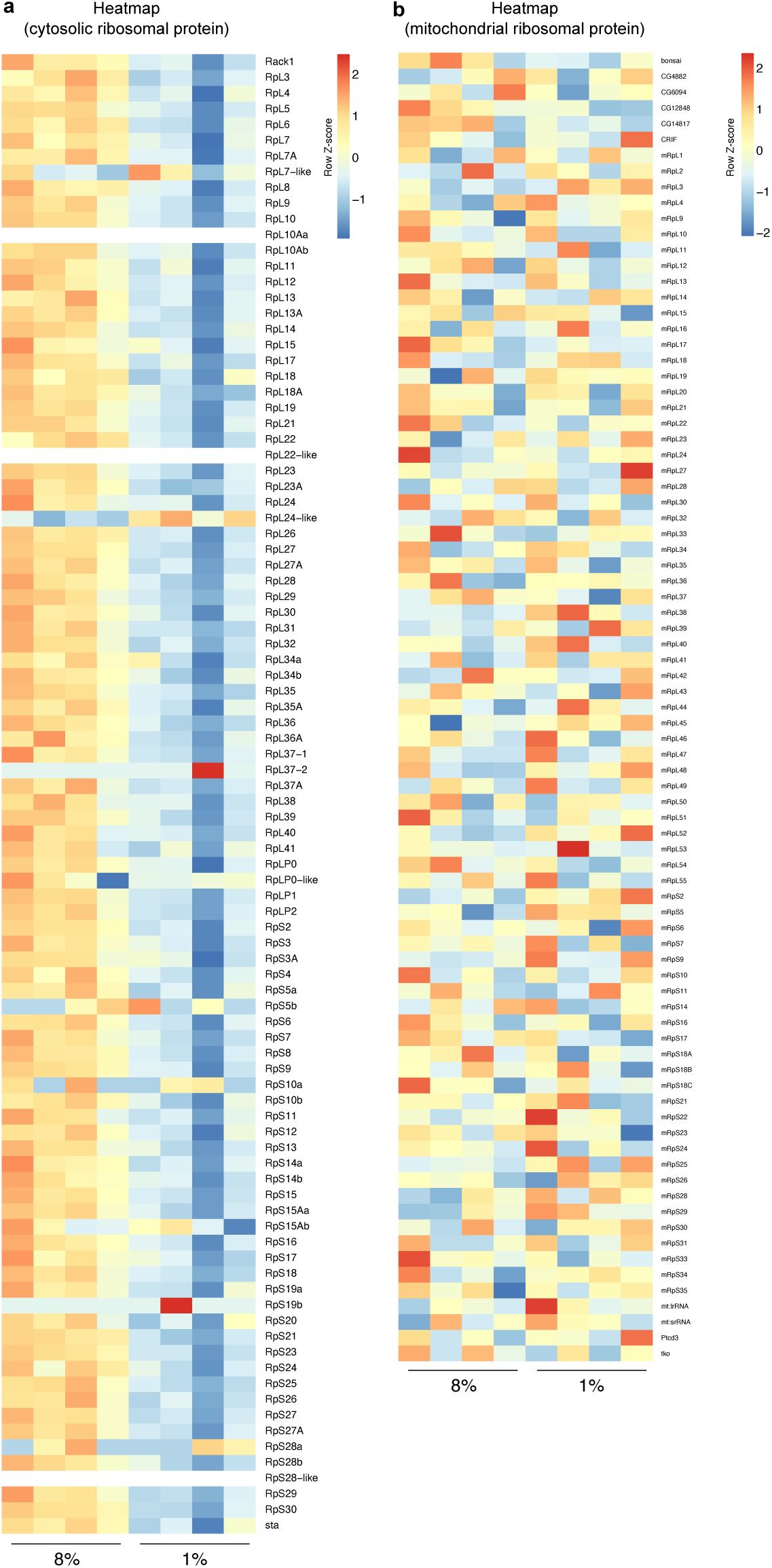
Expression of ribosomal protein encoding genes in adult flies with ePR. **a**,**b**, Heat map of cytosolic (**a**) and mitochondrial (**b**) ribosomal protein encoding genes of Canton-S female flies with ePR at day2.

**Extended Data Fig. 8:**
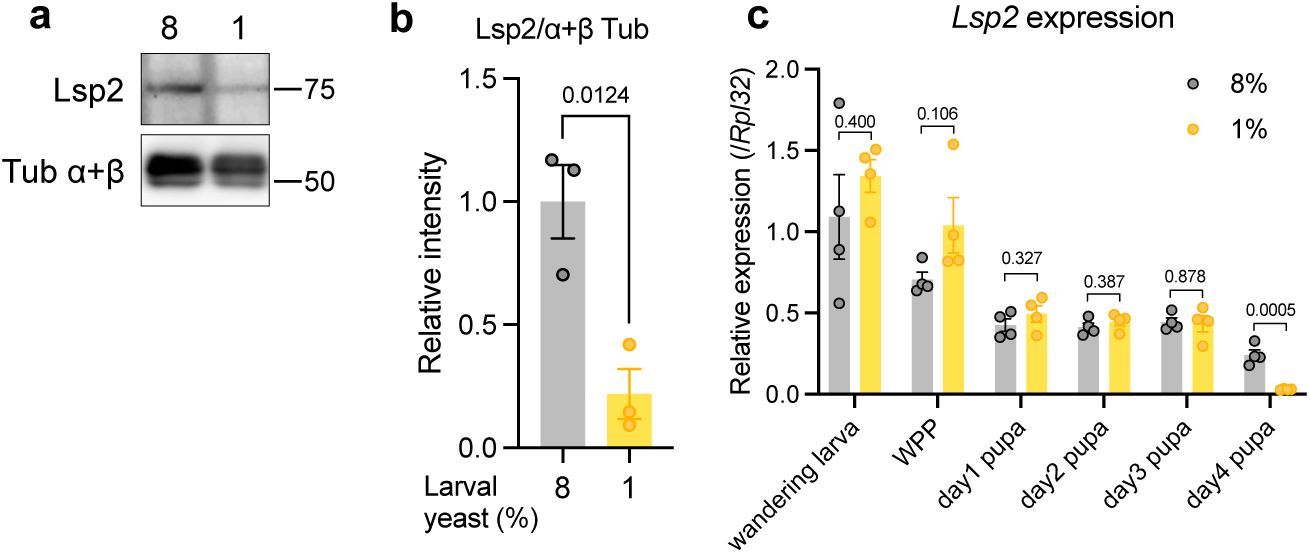
Lsp2 expression level in flies after ePR. **a**,**b**, Representative images of western blotting (**a**) and its quantification (**b**) of the heads of Canton-S female flies with ePR. Antibodies used are anti-Lsp2 and anti-α+β tubulin (loading control). n = 3. **c**, Quantitative RT‒PCR analysis of *Lsp2* in the whole body of Canton-S female flies with ePR. n = 4. For the statistics, a two-tailed Student’s *t* test was used (**b**,**c**). For all graphs, the mean and SEM (**b**,**c**) are shown. Data points indicate biological replicates.

**Extended Data Fig. 9:**
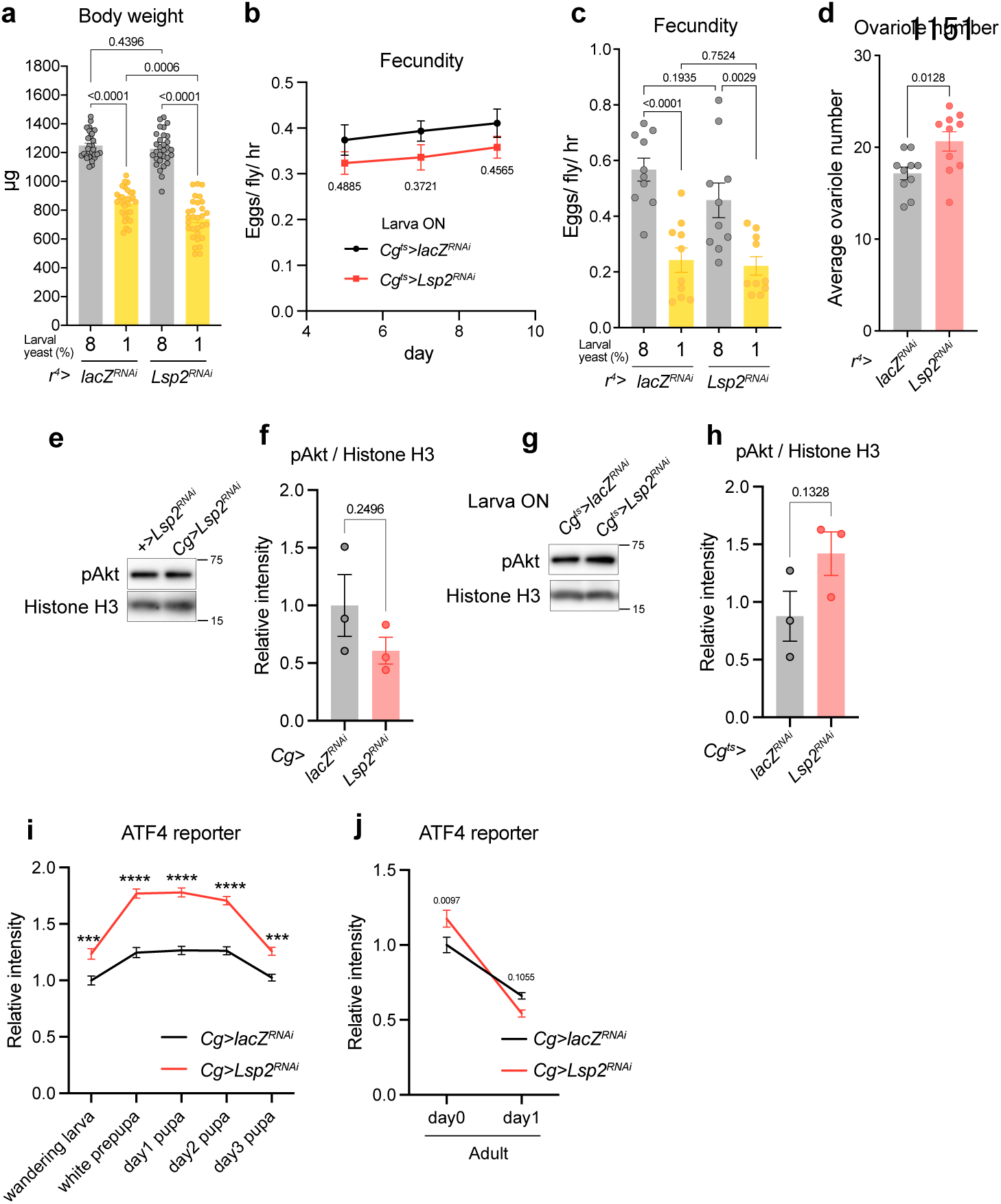
Impacts of *Lsp2* knockdown on body weight, fecundity, and nutritional signalling pathways. **a**, Body weight of female flies with *Lsp2* knocking using fat body driver *r^4^-Gal4* at day 1. n = 30. **b**, Fecundity of female flies from day5 to day9 with *Lsp2* knocking using fat body driver *Cg-Gal4* combined with *tub-Gal80^t^.* Knockdown was performed during larval stage (from 1^st^ instar larva to wandering larva). n = 10. **c**, Fecundity of female flies from day4 to day5 with *Lsp2* knocking using fat body driver *r^4^-Gal4*. Larval diet is ePR. n = 10. **d**, Ovariole number of female flies with *Lsp2* knockdown using *r^4^-Gal4* at day9. n = 10. **e**-**h**, Representative images of western blot analysis (**e**,**g**) and its quantification (**f**,**h**) of adult abdominal carcasses of adult female flies with knockdown using fat body driver *Cg-Gal4* (**e**,**f**) or *Cg-Gal4* combined with *tub-Gal80^ts^*(**g**,**h**). For transient manipulation (**g**,**h**), knockdown was performed during larval stage (from 1^st^ instar larva to wandering larva). The age of flies are day2 (**e**,**f**) or day6 (**g**,**h**). Antibodies used are anti-phosphorylated Akt (pAkt) and anti-histone H3 (loading control). n = 3. **i**,**j**, Quantification of ATF4 reporter (*4E-BP^intron^-dsRed*) fluorescence of whole larvae and pupae (**i**) or adult abdomen (**j**) with *Lsp2* knockdown using fat body driver (*Cg-Gal4*). n = 10. For the statistics, one-way ANOVA with Holm-Šídák’s multiple comparison test (**a**,**c**), two-way ANOVA with Šídák’s multiple comparison test (**b**,**i**,**j**), a two-tailed Student’s *t* test (**d**,**f**,**h**), or was used. \*\*\**P* < 0.001; \*\*\*\**P* < 0.0001. For all graphs, the mean and SEM are shown. Data points indicate biological replicates.

**Extended Data Fig. 10:**
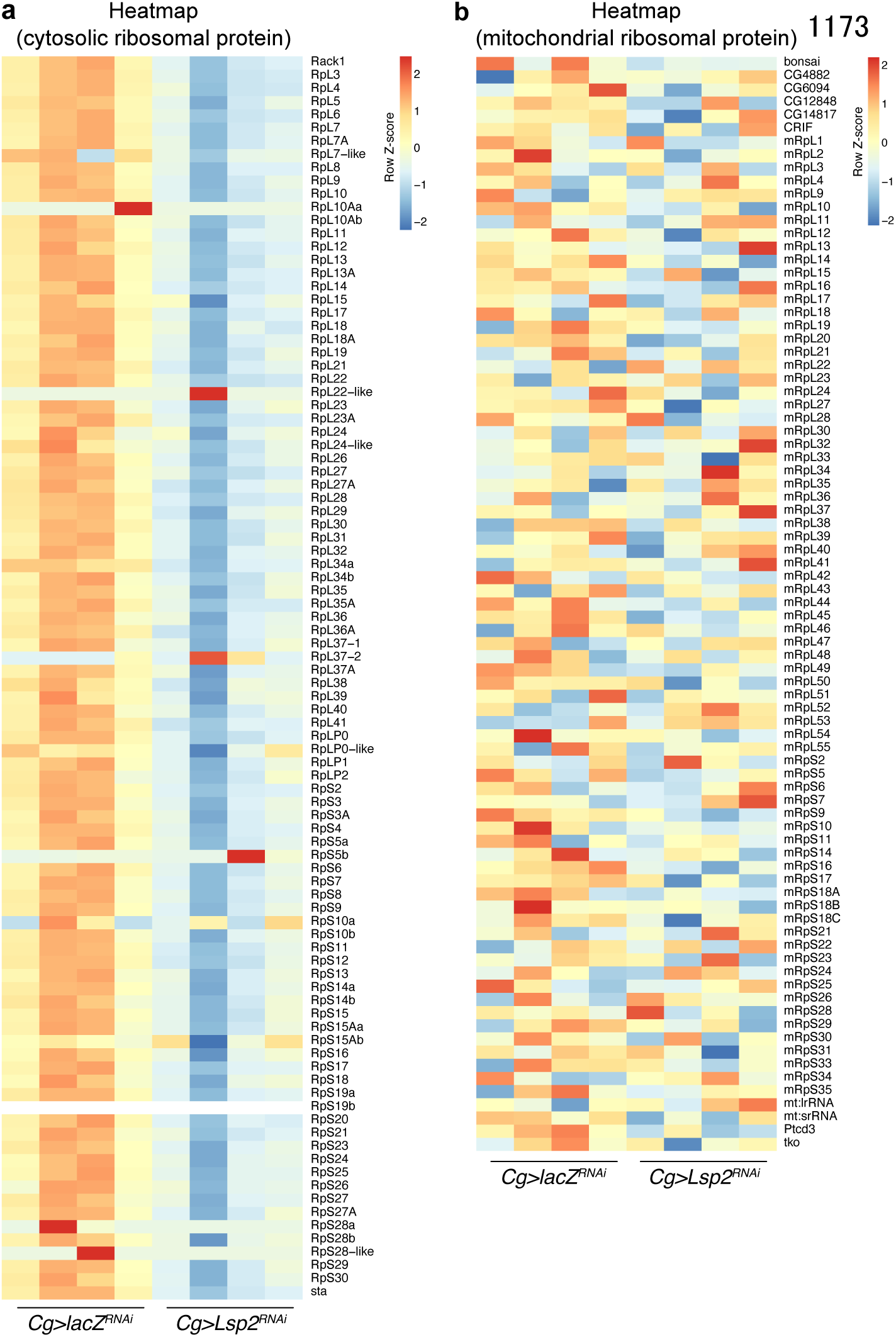
Transcriptome responses of *Lsp2*-knockdown flies. **a**,**b**, Heat map of u cytosolic (**a**) and mitochondrial (**b**) ribosomal protein encoding genes in the head of *Lsp2*-knocked down flies using fat body driver *Cg-Gal4*.

**Extended Data Fig. 11:**
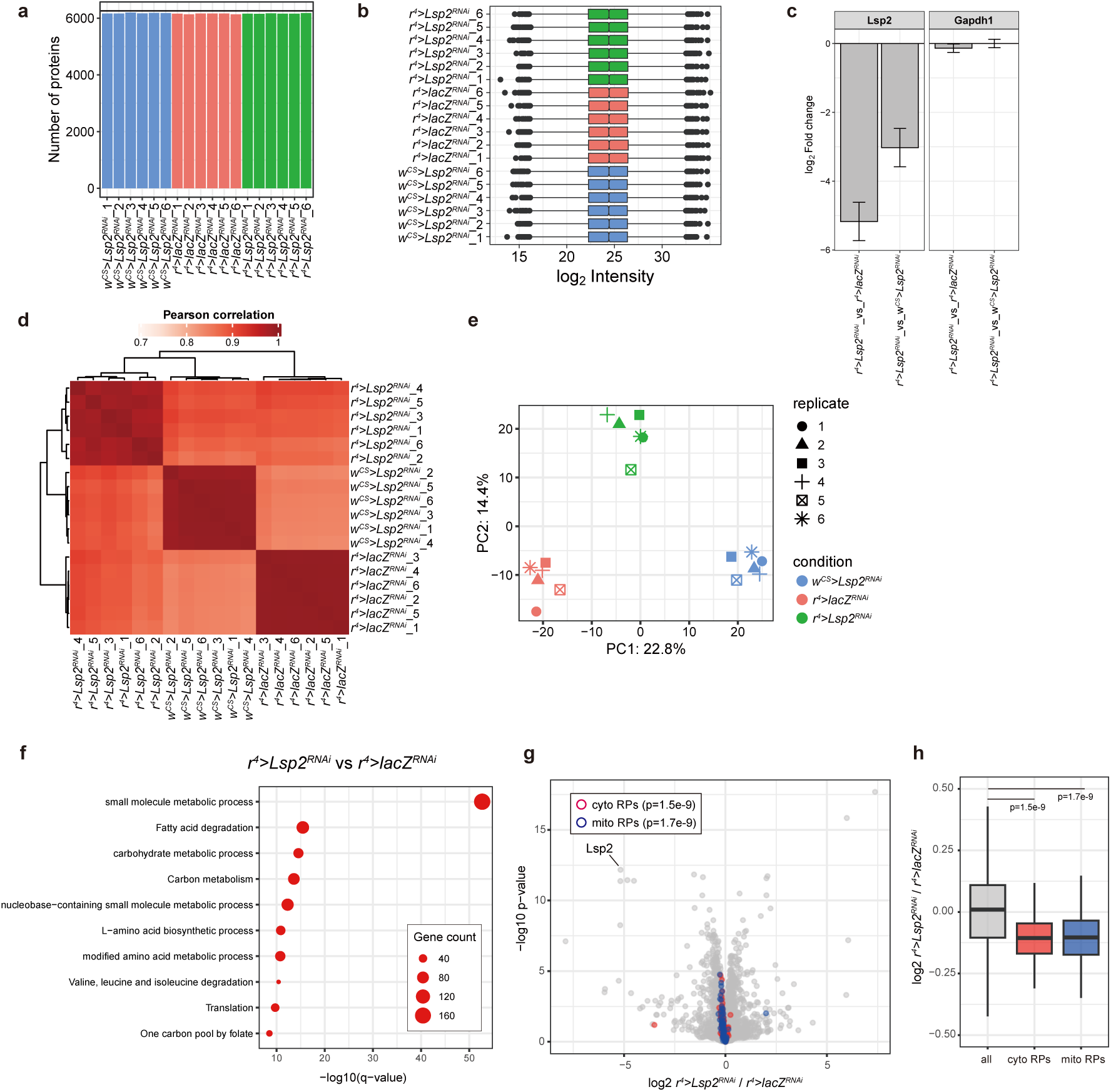
Proteome analysis of Lsp2-knockdown flies. **a**, The number of proteins quantified in each sample: Of the 6,778 proteins identified in total, 6,258 proteins that were consistently quantified across all six biological replicates in at least one condition were included in the analysis. **b**, Overview of log2 protein intensity of individual samples. **c**, Protein abundance of Lsp2 and Gapdh1 (control) with Lsp2-knockdown using fat body driver *r^4^-Gal4*. **d**, Correlation matrix showing Pearson correlation of log2 protein intensity. **e**, PCA plot of individual samples. **f**, GO enrichment analysis of proteins which expression was significantly decreased by Lsp2-knockdown in heads of female Canton-S flies at day2 (p<0.01 and log2 *r^4^>Lsp2^RNAi^*/ *r^4^>lacZ^RNAi^* <-0.1). **g**, A volcano plot showing mean log2 fold change (*r^4^>Lsp2^RNAi^*/ *r^4^>lacZ^RNAi^*) and −log10 p-value of proteins in the head of female flies at day2. The cytosolic and mitochondrial ribosomal proteins are indicated by red and blue circles, respectively. **h**, A box plot showing log2 fold change (*r^4^>Lsp2^RNAi^*/ *r^4^>lacZ^RNAi^*) of proteins grouped into three categories: all proteins, cytosolic and mitochondrial ribosomal proteins. For the statistics, two-sided Wilcoxon rank-sum test (**h**) was used. For the graph, the minimum, the lower quartile, the median, the upper quartile, and the maximum points (**h**) are shown. Data points indicate biological replicates.

**Extended Data Fig. 12:**
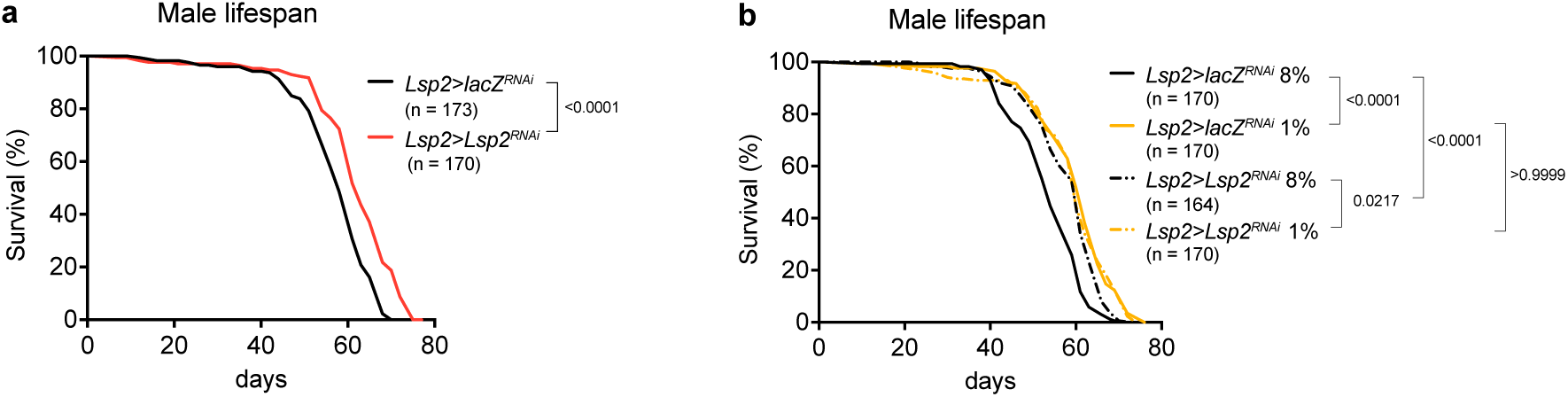
Male lifespan of *Lsp2*-knockdown flies. **a**, Male lifespan of *Lsp2*-knockdown flies using fat body driver *Lsp2-Gal4*. Sample sizes (n) are shown in the figure. **b**, Male lifespan of *Lsp2*-knockdown flies using fat body driver *Lsp2-Gal4* with ePR. Sample sizes (n) are shown in the figure. For the statistics, a log-rank test (**a**,**b**) was used.

**Extended Data Table 1.**
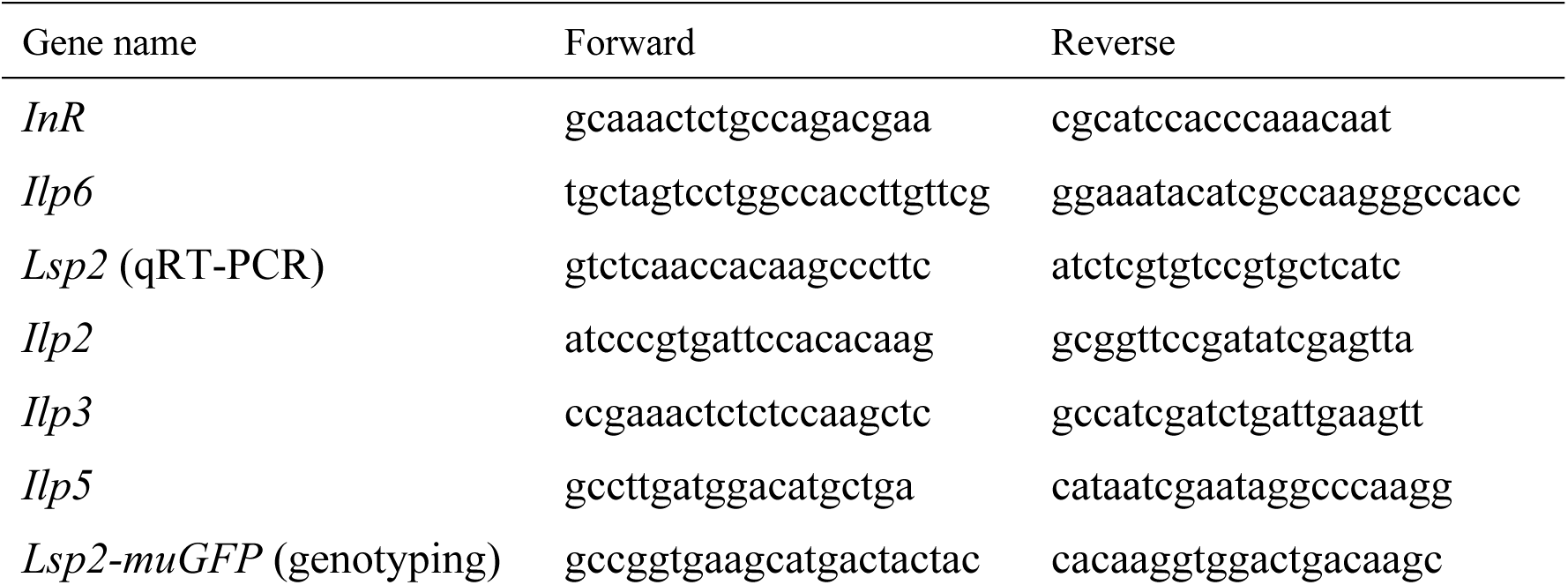
Primer list used in this study for PCR.

